# Predicting Carriers of Ongoing Selective Sweeps Without Knowledge of the Favored Allele

**DOI:** 10.1101/017871

**Authors:** Roy Ronen, Glenn Tesler, Ali Akbari, Shay Zakov, Noah A. Rosenberg, Vineet Bafna

## Abstract

Methods for detecting the genomic signatures of natural selection have been heavily studied, and they have been successful in identifying many selective sweeps. For most of these sweeps, the favored allele remains unknown, making it difficult to distinguish carriers of the sweep from non-carriers. In an ongoing selective sweep, carriers of the favored allele are likely to contain a future most recent common ancestor. Therefore, identifying them may prove useful in predicting the evolutionary trajectory — for example, in contexts involving drug-resistant pathogen strains or cancer subclones. The main contribution of this paper is the development and analysis of a new statistic, the Haplotype Allele Frequency (HAF) score. The HAF score, assigned to individual haplotypes in a sample, naturally captures many of the properties shared by haplotypes carrying a favored allele. We provide a theoretical framework for computing expected HAF scores under different evolutionary scenarios, and we validate the theoretical predictions with simulations. As an application of HAF score computations, we develop an algorithm (PreCIOSS: Predicting Carriers of Ongoing Selective Sweeps) to identify carriers of the favored allele in selective sweeps, and we demonstrate its power on simulations of both hard and soft sweeps, as well as on data from well-known sweeps in human populations.

**Author summary:** Methods for detecting the genomic signatures of natural selection have been heavily studied, and they have been successful in identifying genomic regions under positive selection. However, methods that detect positive selective sweeps do not typically identify the favored allele, or even the haplotypes carrying the favored allele. The main contribution of this paper is the development and analysis of a new statistic (the HAF score), assigned to individual haplotypes. Using both theoretical analyses and simulations, we describe how the HAF scores differ for carriers and non-carriers of the favored allele, and how they change dynamically during a selective sweep. We also develop an algorithm, PreCIOSS, for separating carriers and non-carriers. Our tool has broad applicability as carriers of the favored allele are likely to contain a future most recent common ancestor. Therefore, identifying them may prove useful in predicting the evolutionary trajectory — for example, in contexts involving drug-resistant pathogen strains or cancer subclones.

## Introduction

With genome sequencing, we now have an opportunity to more completely sample genetic diversity in human populations, and probe deeper for signatures of adaptive evolution [1–3]. Genetic data from diverse human populations in recent years have revealed a multitude of genomic regions believed to be evolving under recent positive selection [4–16].

Methods for detecting selective sweeps from DNA sequences have examined a variety of signatures, including patterns represented in variant allele frequencies as well as in haplotype structure. Initially, the problem of detecting selective sweeps was approached primarily by considering variant allele frequencies, exploiting the shift in frequency at ‘hitchhiking’ sites linked to a favored allele relative to non-hitchhiking sites [17,18]. The site frequency spectrum (SFS) within and across populations is often used as a basis for such inference [4,6,19–25]. More recently, methods based on haplotype structure have been developed using a variety of approaches, including the frequency of the most common haplotype [26], the number and diversity of distinct haplotypes [27], the haplotype frequency spectrum [28], and the popular approach of long-range haplotype homozygosity [29–32].

In general, haplotype-based methods seek to characterize the population with summary statistics that capture the frequency and length of different haplotypes. However, the haplotypes are related through a genealogy, and relationships among them are inherently lost in such analyses. In addition, data on the site frequency spectrum can be lost or hidden in analyses focused on haplotype spectra. In this paper, we connect related measures of haplotype frequencies and the site frequency spectrum by merging information describing haplotype relationships with variant allele frequencies. Our main contribution is a statistic that we term the *haplotype allele frequency* (HAF) score, which captures many of the properties shared by haplotypes carrying a favored allele.

Consider a sample of haplotypes in a genomic region. We assume that all sites are biallelic, and at each site, we denote ancestral alleles by 0 and derived alleles by 1. We also assume that all sites are polymorphic in the sample. The *HAF vector* of a haplotype *h*, denoted *c*, is obtained by taking the binary haplotype vector and replacing non-zero entries (derived alleles carried by the haplotype) with their respective frequencies in the sample (Fig. 1A). For parameter 𝓁, we define the 𝓁-HAF *score* of *c* as:

**Figure 1.**
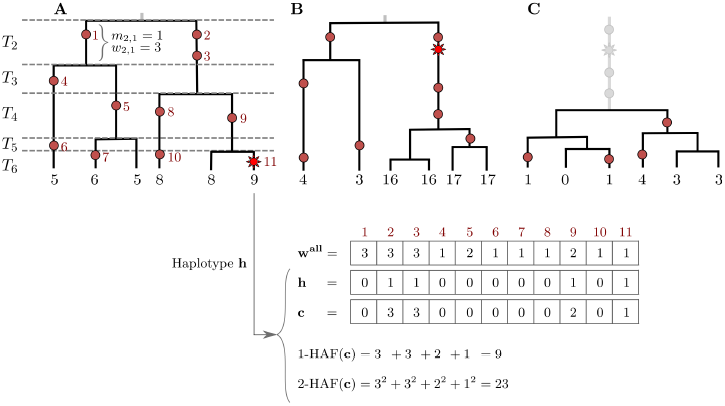
The HAF score. Genealogies of three samples (*n* = 6) progressing through a selective sweep, from left to right. Neutral mutations are shown as red circles, and are numbered in red; the favored allele is shown as a red star. The 1-HAF score of each haplotype is shown below its corresponding leaf, in black. For the rightmost haplotype in (A), the binary haplotype vector **h** is shown along with its HAF-vector c, and 1-HAF and 2-HAF scores. Vector **w** ^al1^ lists the frequencies of all mutations. (A) The favored allele appears on a single haplotype. At this point in time, both the genealogy and the HAF score distribution are largely neutral. Coalescence times (*T*_2_, … , *T*_6_) are shown on the left, where *T*_*k*_ spans the epoch with exactly *k* lineages. (B) Carriers of the favored allele are distinguished by high HAF scores (in large part due to the long branch of high-frequency hitchhiking variation); non-carriers have low HAF scores. (C) After fixation, there is a sharp loss of diversity causing low HAF scores across the sample.

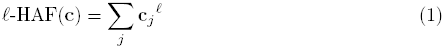

where the sum proceeds over all segregating sites *j* in the genomic region. The 1-HAF score of a haplotype amounts to the sum of frequencies of all derived alleles carried by the haplotype. The 𝓁-HAF score is equivalent to the 𝓁-norm of c raised to the 𝓁^th^ power, or (||c||_𝓁_)𝓁. We will show that during a selective sweep, the HAF score of a haplotype serves as a proxy to its relative fitness.

### Selective sweeps

The classical model for selection, and the one that has received most attention, is the “hard sweep” model, in which a single mutation conveys higher fitness immediately upon occurrence, and rapidly rises in frequency, eventually reaching fixation [17,33]. Under this model, we can partition the haplotypes into carriers of the favored allele, and non-carriers. In the absence of recombination, the favored haplotypes form a single clade in the genealogy. As a sweep progresses, HAF scores in the favored clade will rise due to the increasing frequencies of alleles hitchhiking along with the favored allele. HAF scores of non-carrier haplotypes will decrease, as many of the derived alleles they carry become rare (Fig. 1B). After fixation of the favored and hitchhiking alleles, HAF scores will decline sharply (Fig. 1C), as the selected site and other linked sites are no longer polymorphic. Thus, this reduction in the HAF score results from the sudden loss of many high-frequency derived alleles from the pool of segregating sites [18,20,24]. Finally, as the site-frequency spectrum recovers to its neutral state due to new mutations and drift [23], so will the HAF scores.

Recombination is a source of ‘noise’ for the properties of the HAF score, predicted under the assumption of a hard sweep and no recombination, as it allows haplotypes to cross *into* and *out of* the favored clade. Recombination can lead to (i) haplotypes that carry the favored allele but little of the hitchhiking variation, thus having relatively low HAF scores despite their high fitness, or (ii) haplotypes that do not carry the favored allele but do carry much of the hitchhiking variation, thus having relatively high HAF scores despite their low fitness. By the same logic, recombination adds ‘noise’ after fixation by making the otherwise sharp decline in HAF scores more subtle and gradual. This more gradual decline occurs due to recombination weakening the linkage between the favored allele and hitchhiking variants.

Recently, “soft sweeps” have generated significant interest [34–36]. A soft sweep occurs when multiple sets of hitchhiking alleles in a given region increase in frequency, rather than a single favored haplotype. Soft sweeps may take place by one or more of the following mechanisms: (i) selection from standing variation: a neutral segregating mutation, which exists on several haplotypic backgrounds, becomes favored due to a change in the environment; (ii) recurrent mutation: the favored mutation arises several times on different haplotypic backgrounds; or, (iii) multiple adaptations: multiple favored mutations occur on multiple haplotypic backgrounds. Several methods have been developed for detecting soft sweeps [37,38], as well as for distinguishing between soft and hard sweeps [39–41]. In soft sweeps, multiple sets of hitchhiking alleles rise to intermediate frequencies as the favored allele fixes. This could cause the pre-fixation peak and post-fixation trough in HAF scores to be less pronounced and to occur more gradually compared to a hard sweep.

We find (see Results) that the properties of the HAF score remain robust to many soft sweep scenarios. Moreover, the HAF score could potentially be used to detect soft sweeps. However, in this paper, we focus on the foundations, developing theoretical analysis and empirical work that predicts the dynamics of the HAF score. We also develop a single application. Recall that most existing methods for characterizing selective sweeps focus on identifying regions under selection. Here, given a region already identified to be undergoing a selective sweep, we ask if we can accurately predict which haplotypes carry the favored allele, without knowledge of the favored site. Successfully doing so may provide insight into the future evolutionary trajectory of a population. Haplotypes in future generations are more likely to be descended from, and therefore to resemble, extant carriers of a favored allele. This predictive perspective is of particular importance when a sweep is undesirable and measures may be taken to prevent it. For instance, consider a set of tumor haplotypes isolated from single cells, some of which are drug-resistant and therefore favored under drug exposure. Given a genetic sample of the tumor haplotypes, the HAF statistic may be applied to identify the resistant haplotypes — carriers of a favored allele — before they clonally expand and metastasize.

Below, we start with a theoretical explanation of the behavior of the HAF score under different evolutionary scenarios, validating our results using simulation. We then develop an algorithm (PreCIOSS: Predicting Carriers of Ongoing Selective Sweeps) to detect carriers of selective sweeps based on the HAF score. We demonstrate the power of PreCIOSS on simulations of both hard and soft sweeps, as well as on real genetic data from well-known sweeps in human populations. While our theoretical derivations make use of coalescent theory, and explicitly use tree-like genealogies, we note that HAF scores can be computed for any haplotype matrix including those with recombination events. Our results on simulated and real data imply that the utility of the HAF score extends to cases with recombination as well as other evolutionary scenarios.

## Results

### Theoretical and empirical modeling of HAF scores

We consider a sample of *n* haploid individuals chosen at random from a larger population of size *N*. Let *μ* denote the mutation rate per generation per nucleotide, and let *θ*= 2*NμL* denote the population-scaled mutation rate in a region of length *L* bp. We consider both constant-sized and exponentially growing populations. For exponentially growing populations, let *N*_*0*_ denote the final population size, let *r* denote the growth rate per generation, and let *α* = 2 *N*_*0*_ *r* the population-scaled growth rate. Let *ρ* denote the population-scaled recombination rate. In our theoretical calculations, we assume no recombination (*ρ* = 0), and we derive expressions for the general 𝓁-HAF score. We use simulations to demonstrate the concordance of theoretical and empirical values of the 𝓁-HAF score, and show that the values are robust to the presence of recombination (see ‘Simulations’ in Methods for parameter choices). Although some of our theoretical calculations below derive expressions for the general 𝓁-HAF score, we primarily use 1-HAF in the applied sections. Applications of 𝓁-HAF with 𝓁 > 1 will be explored in future work.

#### Expected 𝓁-HAF score under neutrality, constant population size

First, we assume that the genomic region of interest is evolving neutrally, the population size remains constant at *N*, and that the ancestral states are known or can be derived. In a sample of size *n*, let **c**(*v*) denote the HAF vector c for the *v*^th^ haplotype (*v* ∊ {1,…, n}). Let ξ_*w*_ be the number of sites with derived allele frequency *w*. We only consider polymorphic sites in the sample, so the frequency is in the range *w* ∊ {1,…, *n* − 1}; a mutation present in all or none of the haplotypes in the sample would not be detectable. Each of the ξ_*w*_ sites of frequency *w* contributes *w*^𝓁^ to the 𝓁-HAF score of each of the *w* haplotypes with the mutation, and contributes 0^𝓁^ = 0 for each of the other *n* − *w* haplotypes. The mean of the 𝓁-HAF scores of all *n* haplotypes in the sample is

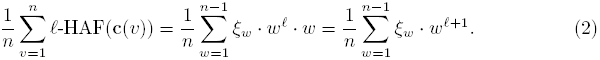

Under the coalescent model, [42, Eq. (22)] shows that 𝔼[ξ_*w*_] = 0/*w* for all 1 ≤ *w* ≤ *n* − 1. By averaging over all haplotypes in all genealogies, the expected 𝓁-HAF score is computed as

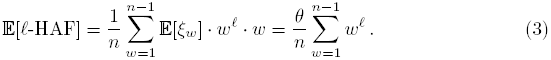

The first two cases (𝓁 = 1, 2) yield

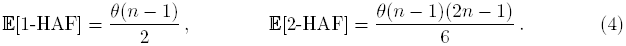

#### Expected 𝓁-HAF score, variable population size

Our derivation of expected 𝓁-HAF scores for constant, neutrally evolving populations does not immediately extend to other demographic scenarios. We describe a second approach that separates coalescence times from the genealogy, and we apply it to compute the expected 𝓁-HAF in an exponentially growing population.

For a sample of size *n*, partition the time spanning from the present back to the sample MRCA into *n* − 1 epochs. Let epoch *k* ∊ {2,…, *n*} be the span of time during which the genealogy contains exactly k lineages (Fig. 1). Note that mutations on a given lineage in a given epoch share the same frequency, as they appear in exactly the same leaves. For example, mutations 2 and 3 in Fig. 1A occur on the same lineage in epoch 2, and they share the frequency 3. Consider the path leading from a randomly chosen haplotype back to the sample MRCA. We can write the 𝓁-HAF score of the haplotype as

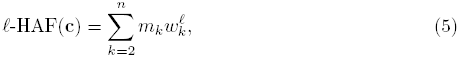

where *m*_*k*_ is the number of mutations that occurred on the path in the *k*^th^ epoch, and *w*_*k*_ is the frequency or weight of those mutations. For a given genealogy with haplotypes *v* ∊ {1,…, *n*}, let **c**(*v*) denote **c** (the HAF vector) for the *v*^th^ haplotype. Similarly, let *m*_*k*_(*v*) and *w*_*k*_(*v*) denote the number of mutations and their frequency in the *k*^th^ epoch for the *v*^th^ haplotype. Epoch *k* splits the haplotypes into *k* equivalence classes, which we call *k-clades.* Let *m*_*k, i*_ and *w*_*k, i*_ denote the corresponding values on the *i*^th^ lineage of the *k*^th^ epoch. We compute the expected value by summing over all haplotypes and genealogies and dividing by *n*. The sum is

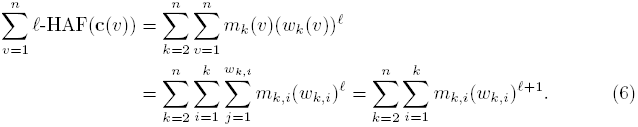

Let *M*_*k, i*_ and *W*_*k, i*_ be random variables denoting the number of mutations, and their frequency respectively, on the *i*^th^ lineage of the *k*^th^ epoch. As the genealogy of a neutrally evolving sample is independent of branch lengths [43], *M*_*k,i*_ and *W*_*k,i*_ are independent random variables. Thus, we can compute the expected 𝓁-HAF score of a randomly chosen haplotype as

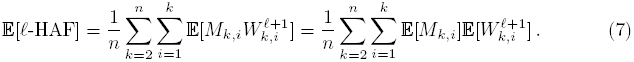

To compute 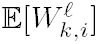, we start with a related quantity. For positive integer 𝓁, denote the *rising factorial*

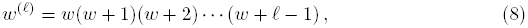

and set *w*^(0)^ = 1. We show in the Appendix that

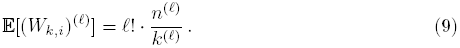

We have *w*^(1)^ = *w* and *w*^(2)^ = *w*(*w* + 1) = *w*^2^ + *w*, so *w*^2^ = *w*^(2)^ − *w*^(1)^, Which leads to:

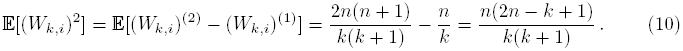

In the Appendix, we generalize this equation to compute 𝔼[(*W*_*k,i*_)^𝓁^]. In addition, we show that for a constant-sized population, the general form in Eq. (7) produces the same result as Eq. (3).

#### Exponential population growth

Eq. (7) can potentially be used to obtain E[𝓁-HAF] under arbitrarily complex demographics. Consider a population of current size *N*_0_ that has been growing exponentially at a rate *r*. The population size at time *t* in the past is given by *N*(*t*) = *N*_0_ *e*^−*rt*^. Exponential population growth is of particular interest, as it has been used to analyze the state of a population under a selective sweep *shortly after fixation.* This is a low point (or *trough*) of observed 𝓁-HAF scores, as early hitchhiking sites have fixed by this time point, and the (relatively recent) sample MRCA is a carrier of the favored mutation. Immediately after fixation, the population — all of which are carriers of the favored allele — has been growing for the duration of the sweep at a rate that is approximately exponential (with growth rate related to the selection coefficient s). In addition, all extant and ancestral haplotypes since the sample MRCA are carriers and therefore equally favored, implying that the independence between *W*_*k,i*_ and *M*_*k, i*_ is kept. While the branch lengths and distribution of *M*_*k, i*_ values change under exponential growth, the distribution for *W*_*k,i*_ remains unchanged as described in Eq. (10). This key insight allows us to use Eq. (7) to estimate the expected HAF scores under exponential population growth.

In order to use 𝔼[*M*_*k,i*_] = *μ*𝔼[*T*_*k*_] under exponential growth, we implement two numerical methods to compute 𝔼[*T*_*k*_]: a ‘cumulative time’ method that uses an approximate distribution of *T*_*k*_ given in [44, p. 559], and a ‘conditional expectation’ method (see Appendix for details). In the conditional expectation method, we compute the expected value of *T*_*k*_ conditioned on *T*_*k*+1_,…, *T*_*n*_, as follows (in the order *k* = *n*, *n* − 1,…, 2):

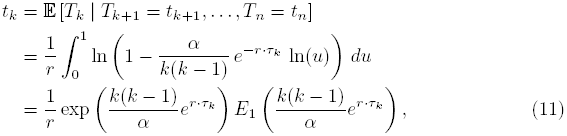

where *α* = 2 *N*_0_ *r* is the scaled growth rate, *τ*_*k*_ = *t*_*k*+1_ + ⋯ + *t*_*n*_ (with *τ*_*n*_ = 0), and *E*_1_(*x*) is the exponential integral 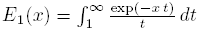

We then use 𝔼[(*W*_*k,i*_)^𝓁^] (evaluated in Eq. (S21)) to evaluate Eq. (7), yielding 𝔼[𝓁-HAF] for exponential population growth as

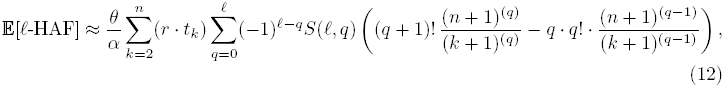

where *S*(*𝓁, q*) denotes the Stirling number of the second kind [45, Ch. 6.1]. We describe these procedures fully in the Appendix.

#### Empirical validation of expected HAF score computation

We tested our theoretical calculations against empirical observations of HAF scores using simulations for neutral evolution with constant population size *N* = 2000 (lowered for subsequent forward simulations; see ‘Simulations’ in Methods). For example, for *θ* = 48 and *n* = 200, the expected 1-HAF is exactly 4776.0 (Eqs. (3),(7)), whereas the empirically observed mean 1-HAF score of 20,000 simulated samples is 4777.1 (Supplementary Fig. S1).

We also modeled exponential growth in population size using the scaled growth rate *α*, using the conditional expectation method in Eq. (12) (see Appendix). As expected, the HAF score is much lower than for constant population size. For *α* = 80, *θ* = 48, *n* = 200, the theoretical mean 1-HAF score is 126.9, whereas the empirical mean of 20,000 simulations is 128.1 (Fig. S2).

We compared the simulations with theoretical expected 𝓁-HAF scores for multiple values of 𝓁 ∊ {1, 2, 3, 4}, and different choices of the population-genetic parameters (*n* ∊ {100, 200, 300,400}, scaled mutation rate *θ* ∊ {24,48}, scaled recombination rate *ρ* ∊ {0, 7.568,15}, and scaled growth rate *α* ∊ {0, 30, 60, 80}). Note that theoretical expected values were computed assuming scaled recombination rate *ρ* = 0, while simulations included *ρ* ∊ {0, 7.568,15}. See Methods for a detailed description of the choice of simulation parameters. Fig. S3 shows the concordance between theoretical and empirical means for each choice of parameters. The concordance improves slightly for increasing values of *n*, 𝓁. Surprisingly, the values are robust to choice of *ρ*, and the variance even reduces slightly for higher *ρ* (Supplementary Fig. S3D).

### HAF score dynamics in selective sweeps

We now consider the dynamics of HAF scores in a population undergoing a selective sweep. To do this, we use data simulated under several scenarios including both hard and soft sweeps. Though soft sweeps can arise under different circumstances, in what follows we restrict our attention to soft sweeps from the standing variation. Fig. 2 illustrates the HAF score dynamics in a single simulated population undergoing a soft sweep, due to standing variation, with selection coefficient *s* = 0.05 and favored allele frequency *v*_0_ = 0.3 when selection begins. See ‘Simulations’ in Methods for a detailed description of the simulation parameters. Initially (leftmost, time 0) the HAF scores of carriers and non-carriers of the favored allele are similar. As the sweep progresses (times 50−200), carrier HAF scores increase to a peak value (HAF-peak). Soon after fixation (time ∼400), we observe a sharp decline in HAF scores (HAF-trough), followed by slow and steady recovery due to new mutation and drift (times 500−4000).

**Figure 2.**
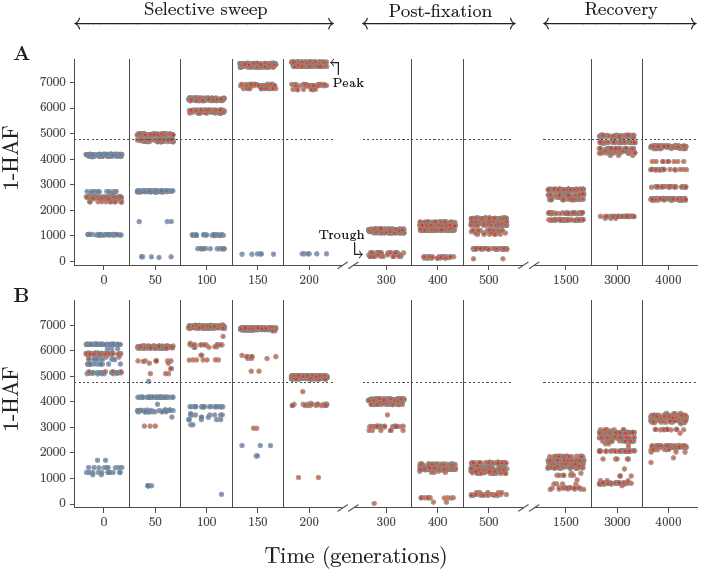
HAF scores in a soft sweep. We consider HAF scores in 50 kb segments, examining *n* = 200 haplotypes sampled from a constant-sized (*N* = 2000) population, evolving with population-scaled mutation rate *θ* = 48 and selection coefficient *s* = 0.05. The soft sweep is modeled by the frequency of the favored allele at the onset of selection, which we set to *v*_0_ = 0.3. We do forward simulations, with time *t* = 0 at the onset of selection and *t* increasing towards the present time. Snapshots of generations are shown at specific times indicated at tick marks on the *x*-axis. Note that these times are increasing but neither consecutive nor regularly spaced. Each selected generation is depicted as a tall thin rectangle. A few rectangles are shown for each phase of a simulated population undergoing a selective sweep. Each point within a rectangle represents the 1-HAF score of a randomly chosen haplotype. Red points represent carriers of the favored allele and blue points represent non-carriers. Points are scattered randomly on the *x*-axis within each rectangle, but all points within the same rectangle represent the same generation at the time indicated by the tick mark on the *x*-axis, regardless of their horizontal position within the rectangle. (A) Simulation of a non-recombining segment. (B) Simulation with population-scaled recombination rate *ρ* = 7.568; see Eq. (17)

Below, we provide a theoretical description of these dynamics, as well as empirical validation using simulations. This allows us to predict HAF scores in (a) the postfixation trough; (b) the pre-fixation peak; and (c) the rate of growth of HAF scores from pre-sweep to peak value.

#### Empirical validation of the post-fixation HAF-trough

We showed using simulations that the HAF score computations for an exponentially growing population (Eq. (12)) also approximate a population evolving under a selective sweep *shortly after fixation.* This enables prediction of the HAF-trough value.

The HAF-trough of a sweep is the value of 1-HAF at fixation. We took the mean of the HAF-trough values over 200 populations simulated under selective sweeps with coefficients *s* ∊ [0.02,0.08] (see ‘Simulations’ in Methods), and compared it to 1-HAF values in simulated neutral populations growing exponentially at rates *α* ∊ [32,101]. Fig. 3A shows a close similarity between the 1-HAF values under exponential growth (blue) and the selective sweep trough (red). The resemblance is closer for higher values of *s* and *α*. To some extent, this may stem from the increased simulation variance we see for lower values of *s*. On a deeper level, selective sweeps with fixed population size demonstrate a logistic curve [46, Sec. 6.1.3]. However, for higher values of *s* and *α*, the concordance between selective sweeps and exponential growth is increased.

**Figure 3.**
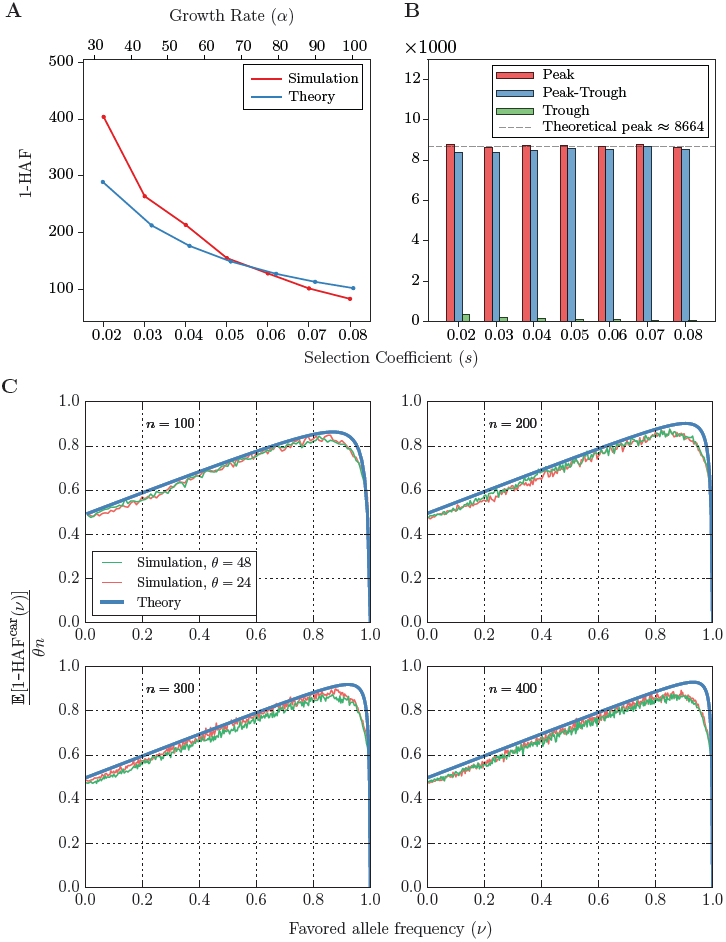
HAF scores in a selective sweep ‘peak’ and ‘trough’. (A) Observed values (red) of the mean ‘trough’ 1-HAF scores in simulated selective sweeps with coefficients *s* ∊ [0.02, 0.08] and theoretical values (blue) of expected 1-HAF scores under exponential population growth with population-scaled rates *α* ∊ [32,101] given by Eq. (12). Simulated 1-HAF scores (red) represent the mean of 200 simulated population samples for each value of *s*, with *θ* = 48, *n* = 200. (B) Observed mean 1-HAF peak, trough, and difference (peak–trough) for selective sweeps with coefficients *s* ∊ [0.02,0.08]. The dashed line represents the approximate value of the peak 1-HAF score given by Eq. (15). (C) Dynamics of the expected value of 1-HAF^car^ (1-HAF score of haplotypes carrying the favored allele) plotted as a function of the fraction of carriers (*v*) in the sample during a selective sweep. For each (*θ*, *n*, *v*) with *θ* ∊ {24, 48}, *n* ∊ {100, 200, 300,400}, 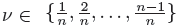, *s* = 0.08, and *N* = 2000, we plotted the mean value of (1-HAF^car^)/(*θn*) over 1500 trials, and compared against the theoretical values (Eq. (13)).

#### The pre-fixation 1-HAF-peak

As the selective sweep progresses, the value of the HAF score of haplotypes carrying the favored allele increases, eventually reaching a peak value. Consider *n* haplotypes sampled from a fixed population of *N* haploid individuals under a selective sweep. Let *μ* denote the mutation rate per base per generation in the genomic region of interest, and assume that there is no recombination. The scaled mutation rate is given by *θ* = 2*Nμ*.

We let *v* denote the fraction of carrier haplotypes in the sample. When *v* ≤ 1/n (i.e., 0 or 1 carriers), there is no selection going back in time, and the time to MRCA can be computed using the neutral Wright-Fisher model [47]. The expected 1-HAF scores for carriers and non-carriers are identical (Eq. (3)). At the time when *v* first equals 1, there are no non-carriers, and the HAF-scores are given by the exponential growth model. In the Appendix, we model the 1-HAF scores for all intermediate values of *v*.

Let 1-HAF^car^ (respectively, 1-HAF^non^) denote the 1-HAF score of a random haplotype carrying the favored allele (respectively, a non-carrier) when a fraction *v* of the *n* sampled haplotypes carry the favored allele. In the Appendix, we show that under strong selection (*N*_*s*_ > > 1) and no recombination (*ρ* = 0),

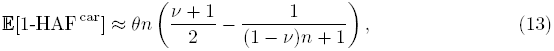

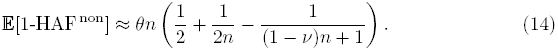

For any sample of size *n*, the carrier haplotypes reach a peak value of 1-HAF^car^ as v varies along its trajectory. We do not compute the expected value of this peak (𝔼[max_*v*_(1-HAF^car^(*v,n*))]) directly. Instead, we compute the peak value of 𝔼[1-HAF^car^(*v, n*)] (maximizing over all *v* ∊ [0,1]) as

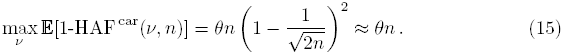

Note that under strong selection, this peak does not depend on *s* (see Fig. 3B).

The trough for each trajectory is computed as the 1-HAF score at fixation (when *v* = 1 is first reached).

#### Empirical validation

Simulated data under selective sweeps with coefficients *s* ∊ [0.02, 0.08] show that (i) the pre-fixation HAF peak scores appear to be independent of the selection coefficient (Fig. 3B), and (ii) as predicted by Eq. (15), the mean value of the HAF peak score is approximately *θn* (Fig. S10 in Appendix). We also simulated (1-HAF^car^)/(*nθ*) as a function of *v* (Fig. 3C and Fig. S9). The results show a tight correspondence between theory and empirical observations.

### HAF score Application: Characterizing carriers and non-carriers

Our understanding of the dynamics of HAF scores of a haplotype during a selective sweep has many potential applications. For example, we could compare the dynamics of hard and soft sweeps to distinguish between the two events. Second, conditioning on known or deduced selective sweeps in a population sample, we can predict the state (carrier/non-carrier) of the favored allele in its haplotypes. Below we explore the latter, leaving the former to future work.

In Fig. 4A we show the distributions of haplotype 1-HAF scores aggregated from 500 simulated populations undergoing a hard selective sweep (see ‘Simulations’ in Methods for detailed parameter choices). Scores were computed for random samples of *n* = 200 haplotypes taken at regular time intervals. They are stratified by the frequency of the favored allele at the time of sampling. Further, scores are stratified into carrier and non-carrier classes (of the favored allele). As with a single population, HAF scores of carriers and non-carriers diverge as the sweep progresses in frequency. We note, however, that even close to fixation (frequencies 80−100%) the distributions of HAF scores between carriers and non-carriers maintain considerable overlap. The high variance in HAF scores makes them only weakly informative of sweep carrier status when comparing across population samples (or genomic regions within a single population). Within a single population sample, however, the HAF scores are highly informative of the carrier status. This is illustrated in Fig. 4B, showing the distributions of HAF score percentile rank within their respective samples. We observe that the rank distributions have minimal overlap for carriers and non-carriers of the favored allele. Any remaining overlap in the percentile rank distributions in the final stages of a sweep (favored allele frequency ≥70%) stems mostly from recombination, which allows the favored allele to recombine onto haplotypes outside the selected clade (creating low HAF score carriers) and vice-versa (creating high HAF score non-carriers). The overall strong separation between carriers and non-carriers is further illustrated by the highly significant *P*-values of Wilcoxon rank sum tests rejecting the null hypothesis of identically distributed HAF scores among carriers and non-carriers *within* each population sample (Fig. 4C).

Fig. 4 does not show how HAF scores are distributed following fixation of the sweep. Starting at fixation, we see a strong decline in HAF scores owing to the loss of many high frequency derived alleles from the pool of segregating sites. However, crossover events may unlink hitchhiking alleles from the favored allele, and they may remain segregating in the population even after fixation of the favored allele. Therefore, the decline in HAF scores may be abrupt or gradual, depending on the linkage between the favored and hitchhiking alleles. Finally, after reaching a trough, HAF scores gradually recover to their neutral levels over time. The post-fixation dynamics of HAF scores are shown in Fig. S5.

**Figure 4.**
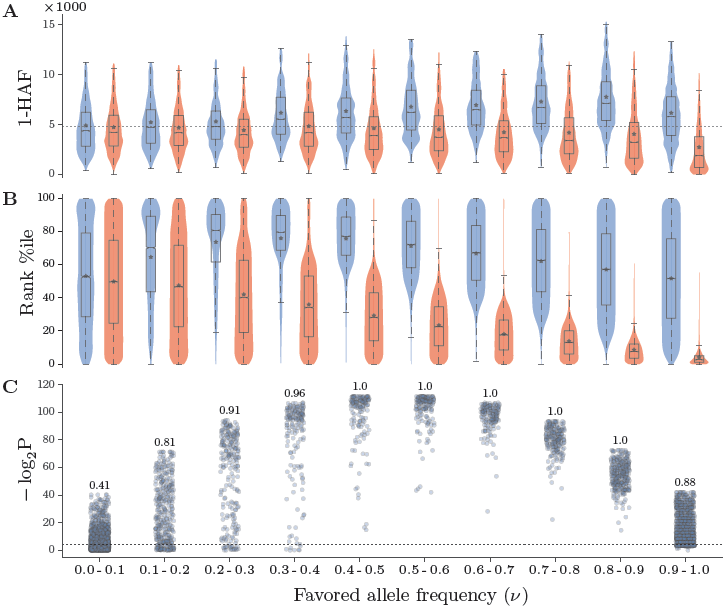
HAF score dynamics in ongoing selective sweeps. HAF scores were computed from 500 simulated population samples (*n* = 200) undergoing a hard sweep (*θ* = 48, *ρ* = 7.568, *s* = 0.05). (A) Each violin shows the Gaussian kernel density estimation (KDE) of 1-HAF scores in carriers (blue) and non-carriers (red) of the favored allele, as the sweep progresses in frequency. A standard box plot is overlaid on each violin (outlier points omitted), and the means are indicated by stars. The horizontal dotted line represents the neutral expectation, as derived. (B) Corresponding violins showing the *in-sample* percentile rank of 1-HAF scores. (C) − log_2_(P) values for Wilcoxon rank sum tests rejecting the null hypothesis of identically distributed 1-HAF scores among carriers and non-carriers *within* each population sample. The number above each bin indicates the fraction of significant tests (where *P* < 0.05, shown by the dotted line).

#### PreCIOSS: <u>Pre</u>dicting <u>C</u>arriers of <u>O</u>ngoing <u>S</u>elective <u>S</u>weeps

Our simulations suggest that, in a region undergoing a selective sweep, we could use HAF scores to predict whether a haplotype is carrying the favored allele. We implemented a simple algorithm (PreCIOSS) to carry out this prediction by clustering HAF scores in a sample. PreCIOSS takes as input a set of binary haplotypes sampled from a population undergoing a selective sweep. For each haplotype, the 𝓁-HAF score is computed (𝓁 = 1 by default). We then fit a Gaussian Mixture Model (GMM) with exactly two Gaussians to the haplotype HAF scores. The fit is performed using Expectation Maximization (EM). Finally, we apply the fitted model to assign a label to each haplotype according to the Guassian component to which it is assigned. Haplotypes whose HAF score is higher are denoted as ‘carriers’.

We apply PreCIOSS to data from simulated populations undergoing hard and soft sweeps (see ‘Simulations’ in Methods). The haplotypes predicted as carriers might in fact be carriers (True Positives, TP) or non-carriers (False Positives, FP). Similarly, the haplotypes predicted as non-carriers could be True Negatives (TN) or False Negatives (FN). We measure the *balanced accuracy*

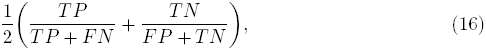

which is more appropriate to use than *Rand accuracy* (*TP* + *TN*)/(total predictions) when the positive and negative classes appear at different proportions in the sample [48].

The balanced accuracy of PreCIOSS on hard sweeps is shown in Fig. 5A for a specific choice of parameters (500 samples with *n* = 200, *θ* = 48, *ρ* = 7.568, *s* = 0.05). Once the sweep reaches frequencies above 30%, the balanced accuracy increases (median ∼70%) and remains high (median ∼90%) for the remainder of the sweep. At the very beginning and end of the sweep, the balanced accuracy, despite being asymptotically unbiased, suffers from high variance due to the severe class imbalance (few carriers in the beginning, few non-carriers at the end). The accuracy is high for soft sweeps (Fig. 5B, run with similar parameters, and *v*_*0*_ = 0.3), where the higher initial carrier frequency makes for a faster increase in accuracy. As expected under soft sweeps, the overall balanced accuracy is somewhat reduced relative to hard sweeps.

**Figure 5.**
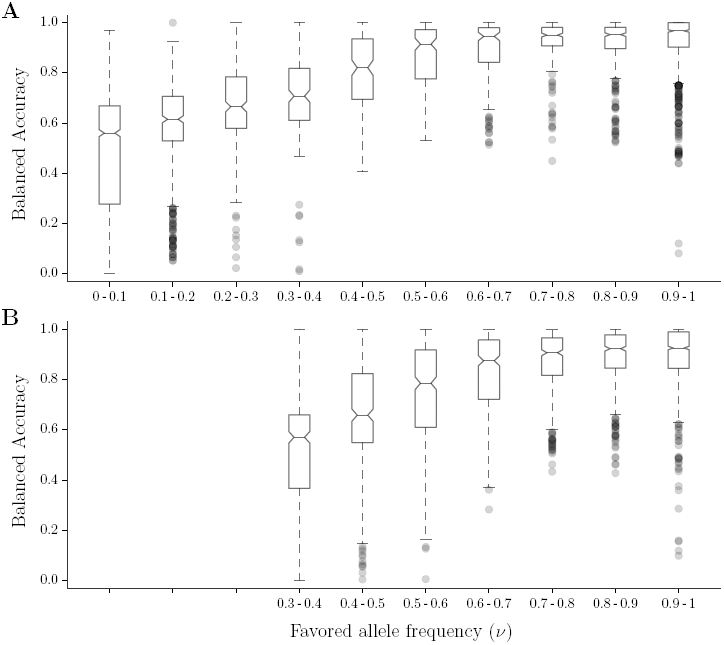
Predicting carriers of hard and soft sweeps. Balanced accuracy (Eq. (16)) of PreCIOSS in populations undergoing hard and soft sweeps. A total of 500 population samples were simulated (*n* = 200, *θ* = 48, *ρ* = 7.568) undergoing (A) a hard sweep (*s* = 0.05, 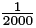), and (B) soft sweep (*s* = 0.05, *v*_*0*_ = 0.3). We split each sweep into intervals as *v* progresses ([0, .1], [.1, .2], etc.). Within each interval for *v*, we show the distribution of balanced accuracies observed using standard box plots.

We tested PreCIOSS under a wide range of population-genetic and sweep parameters (Table S1), and observed consistently high balanced accuracy in carrier-state prediction as the sweeps progress (see Fig. S6). Generally, PreCIOSS has slightly better balanced accuracy on hard sweeps than on soft sweeps. A higher recombination rate has only a limited impact (see Fig. S6A,E), suggesting that PreCIOSS will be robust to different recombination scenarios.

### Applying PreCIOSS to human selective sweeps

To evaluate the effectiveness of PreCIOSS in distinguishing carriers of a selective sweep from non-carriers, we applied it to several genomic regions (e.g., [39]) where (i) there is strong evidence of a selective sweep, and (ii) the favored allele has been characterized. In applying PreCIOSS to the datasets, we assumed that the region was known, but did not supply the favored allele to PreCIOSS. In each case, we tested if PreCIOSS could separate the haplotypes that carried the favored allele. We use phased haplotypes from the HapMap project [49], setting the ancestral allele to that observed in orthologous Chimpanzee sequence [50].

#### LCT

We consider the well-known sweep in the lactase (LCT) gene region in Northern Europeans. The best characterized variant is C/T-13910 (rs4988235), for which the T allele was found to be 100% associated with lactase persistence in the Finnish population [51]. T-13910 was further shown to be causal by in-vitro analysis, where it was found to increase enhancer activity [52,53]. We considered haplotypes from the CEU population, applying PreCIOSS to a 50 kb window centered at C/T-13910 (Fig. 6A). This yielded 100% accuracy in classifying carriers from non-carriers. Increasing the window size above 50 kb, the balanced classification accuracy reduced to ∼90% (Fig. 6B). LCT shows the highest and most prolonged (with increasing distance from the causal site) statistical significance in separating carrier and non-carrier HAF scores, remaining highly significant for haplotypes of 1.5 Mb (Fig. 6B, blue line). Despite this highly significant separation, the classification accuracy is initially unstable, alternating between ∼100% (perfect classification) and 90%. This is due to the pattern of HAF scores observed in the LCT region, where carriers form a tight cluster with the highest scores, but several non-carriers cluster closer to carriers than to the majority of other non-carriers (Fig. 6A). These haplotypes are therefore sometimes included (90% accuracy) and sometimes excluded (100% accuracy) from the reported ‘carriers’ class.

**Figure 6.**
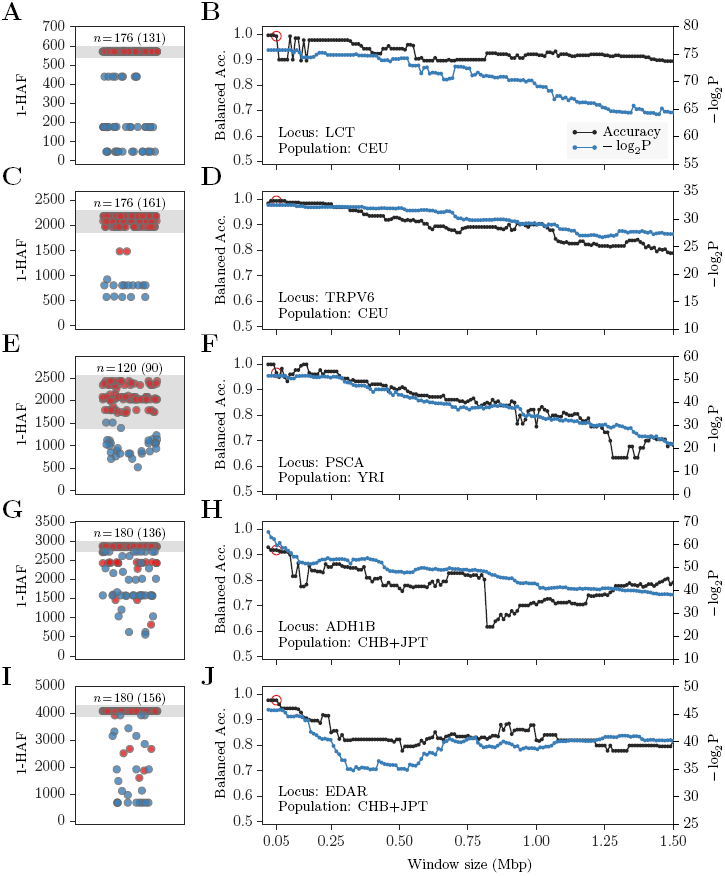
Predicting carriers of well-known selective sweeps. (Left): Haplotype 1-HAF scores in a 50 kb window centered at known favored sites indicated by the gene name, and the SNP identifier. (A) LCT/rs4988235, (C) TRPV/rs4987682, (E) PSCA/rs2294008, (G) ADH1B/rs1229984, and (I) EDAR/rs3827760. Points represent haplotype 1-HAF scores, red indicating a carrier of the favored allele and blue indicating a non-carrier. At the top of each panel, the number of haplotypes, n, is shown, with the number of carriers in parenthesis. Areas shaded in gray indicate haplotypes designated as ‘carrier’ by PreCIOSS. (Right) classification Balanced Accuracy (black) and − log_2_(*P*) values (blue) as function of window size around the favored allele in (B) LCT, (D) TRPV6, (F) PSCA, (H) ADH1B, and (J) EDAR. *P*-values are for Wilcoxon rank sum tests rejecting the null hypothesis of identically distributed 1-HAF scores among carriers and non-carriers. Red circles indicate the 50 kb windows shown on the left.

#### TRPV6

Transient Receptor Potential Cation Channel, Subfamily V, Member 6 (TRPV6) is a membrane calcium channel thought to mediate the rate-limiting step of dietary calcium absorption. It is reportedly under strong positive selection in several non-African populations [54,55]. Following Peter et al. (2012) [39], we focus on the CEU population and set rs4987682 as the favored allele. This site, of the three non-synonymous SNPs with highest allele frequency differentiation among human populations, is the only one located in the the N-terminal region of TRPV6, thought to be the target of selection [56]. Applying PreCIOSS to a 50 kb window centered at this site, we obtain ∼99% balanced classification accuracy (Fig. 6C), which gradually decays to ∼80% when considering a 1.5 Mb window (Fig. 6D). As with LCT, the separation in HAF scores between carriers and non-carriers of the allele is highly statistically significant (P < 10^−7^; see Fig. 6D). Unlike LCT, accuracy decays stably with distance from the favored site. This appears to be due to the less complex clustering pattern of non-carrier HAF scores in the region (Fig. 6C).

#### PSCA

Prostate Stem Cell Antigen (PSCA) has been proposed to be under selection in a global analysis of allele frequency differentiation [57]. The putative causal site is a non-synonymous SNP (rs2294008) known to be involved in several cancer types [58,59]. Interestingly, the derived allele is observed in all human populations, but at vastly different frequencies. It is most common in West Africa and East Asia, where it segregates at ∼75% frequency. We consider haplotypes from the YRI population and apply PreCIOSS to a 50 kb window centered at rs2294008, yielding balanced accuracy of 97% (Fig. 6E). Unlike LCT and TRPV6, accuracy decays more noticeably with distance, reaching 63% at 1.25 Mb (Fig. 6F). This sharper decay in accuracy is even more pronounced when considering the sweep in the CHB population (Fig. S4). Such decay is consistent with a (*soft*) sweep from the standing variation, which would allow more time for recombination to break the linkage between the favored allele and hitchhiking variation. Indeed, the sweep in PSCA was proposed to be from the standing variation by Bhatia et al. (2011) [57], and further substantiated as such by Peter et al. (2012) [39].

#### ADH1B

ADH1B encodes one of four subunits of Alcohol dehydrogenase (ADH1), which plays a key role in alcohol degradation. ADH1 genes (including ADH1B) have been studied extensively on both a functional and a population-genetic level, as they are thought to be one of the major drivers of alcoholism risk [60]. These genes have also been suggested to cause the “alcohol flush” phenotype common in Asian populations [61]. A specific non-synonymous mutation in ADH1B (Arg47His, rs1229984) has been proposed to be the target of selection. This is because (i) the derived allele has been shown to cause increased enzymatic activity [62,63], and (ii) the estimated age of the allele coincides with rice domestication [61,64] and the availability of fermented beverages [65]. Computing HAF scores for phased haplotypes from East Asian populations (CHB+JPT) and applying PreCIOSS, we obtained balanced classification accuracy of 92% using a 50 kb window centered at rs1229984 (Fig. 6G). Both accuracy and statistical significance (of class separation) gradually decay with increasing window size (Fig. 6H). As before, statistical significance decays more stably than classification accuracy.

#### EDAR

EDAR encodes a cell-surface receptor and has been associated with development of distinct hair and teeth morphologies [66,67]. Specifically, a non-synonymous SNP (rs3827760, V370A) has been associated with these phenotypes [68]. The SNP is located within a DEATH-domain, which is highly conserved in mammals [69], and has been experimentally confirmed (in vitro) to increase EDAR activity [68]. It is found at very high frequencies in East Asian and American populations, while being completely absent from Europeans and Africans [68]. The EDAR gene has been found to be under selection in multiple studies [30,54,70], showing one of the strongest signatures of selection genome wide among the 1000 Genomes populations [39]. Applying PreCIOSS to phased CHB+JPT haplotypes in a 50 kb region centered at rs3827760, we obtained 98% balanced accuracy in predicting carriers vs. non-carriers of the allele.

In each case, PreCIOSS was applied to a 50 kb window centered at the favored allele, and separated the carriers and non-carriers with high accuracy of 97-100% (Fig. 6A–F). The accuracy decayed with increasing window size, but in many cases stayed high even for windows of 1.5 Mbp.

## Discussion

This paper introduces a new perspective on the genetic signatures of selective sweeps. From identifying and characterizing sweeps in a population sample — the topic of typical studies of selective sweeps — we progress to considering the role of individual haplotypes within an ongoing sweep. Using both simulated and real data, we show that the HAF score is well-correlated with the *relative* fitness of individual haplotypes, and that our algorithm (PreCIOSS) is highly effective at predicting carriers of selective sweeps.

The HAF framework has many natural extensions and potential applications. On the theoretical side, we have obtained the expected HAF score in both constant-sized and exponentially growing populations evolving neutrally (Eqs. 3,12). However, we do not yet know the variance. This quantity would provide a better understanding of the respective distributions, and a means to to statistically test for deviations from neutrality. Moreover, although we have observed in simulation and in practice that our theoretical argument is robust to recombination (genealogies violating a tree structure), a theoretical argument supporting these observations would be valuable.

In terms of application, several additional directions are worth investigating. The HAF framework is potentially useful in distinguishing hard from soft sweeps. Intuitively, hard sweep genealogies will likely have a single hitchhiking branch dominating the HAF scores, and leading to near-uniform scores in favored haplotypes. However, soft sweep genealogies may have several hitchhiking branches, potentially leading to distinct HAF score peaks. Even if the different favored clades happen to have similar scores, the haplotypes within them will not form a highly-related group as expected in hard sweeps.

Our results on known selective sweeps in humans illustrates this idea already (Fig. 6). A recent study by Peter et al. (2012) [39] assigned posterior probabilities to hard vs. soft sweeps occurring in the same genes. Peter et al. assigned the highest likelihood of a hard sweep to LCT (0.99), followed by EDAR (0.89), ADH1B (0.78), TRPV (0.45), and finally PSCA (0.24). This is in striking concordance with the spread in HAF scores in Fig. 6. The clusters capturing the carriers in LCT and EDAR have tightly distributed HAF scores (Fig. 6A,I). The cluster for ADH1B (Fig. 6G) has more variance by comparison, and the variance increases for TRPV6 (Fig. 6C) and PSCA (Fig. 6E), with PSCA showing the highest variance of HAF scores in carriers.

Finally, perhaps the highest potential impact of the HAF score could be in predicting the ‘MRCA of the future’. We know that future haplotypes are more likely similar to favored individuals than to unfavored ones, and that HAF scores correlate well with relative fitness in ongoing selective sweeps. Therefore, high HAF haplotypes are more likely to be similar to future generations. This relationship is particularly valuable when action may be taken based on such predictions. For instance, rapid influenza viral evolution is known to change the strain composition from year to year. The mutations are a mix of favored and deleterious mutations. The fitness and frequency of the current year‘s strain have been used to predict the next year’s dominant strain [71]. The HAF score may allow for a careful look at the dynamics of the current strain and possibly offer better insight into the problem. As a second example, tumor cells show great heterogeneity and much variation occurs at the single cell level. This intra-tumor variation allows sub-population of cells to resist therapy and proliferate [72]. Once again, HAF scores of haplotypes in cells undergoing treatment can potentially distinguish between carriers and non-carriers of drug resistance mutations, and thereby improve our insight into mechanisms of drug resistance.

## Methods

### Simulations

We simulated data for various evolutionary scenarios. Neutral samples were generated using the coalescent (backwards) simulator *ms* [73]; sweep samples were generated using the Wright-Fisher (forwards) simulator *mpop* [5]. All simulations generated samples of *n* ∊ {100, 200, 300, 400} haplotypes from a larger effective population of *N* = 2000 haplotypes, each of length 50 kb. A mutation rate of approximately *μ* = 2.4 · 10^−8^ mutations per bp per generation was used [74,75]. For simulating populations from a region of length *L* = 50 kb, and assuming effective population size of 10, 000 individuals (20, 000 haplotypes), we obtain *θ* = 2*N μL* = 24. For our simulations, we choose *θ* ∊ {24, 48}. However, we worked with a smaller value of *N* = 2, 000 haplotypes to reduce the computational requirements of forward simulation, and compensated by appropriately scaling the mutation rate to 2.4 · 10^−7^ to maintain the scaled mutation rate, *θ*.

The simulations either had no recombination, or used 3.784 · 10^−8^ crossovers per generation, per base (e.g. [76]). For a 50 kb region, we used the population-scaled rate

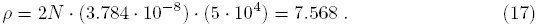

We also worked with a higher population-scaled rate of *ρ* = 15.

For exponential growth, we used *N* = 2000 as the size of the final (current) population. Let *r* denote the growth rate per generation, so that at *t* generations prior to the current generation, the population size was *N*(*t*) = *Ne*^−*rt*^. Define the scaled growth rate *α* = 2*Nr*. We set α to a range of values in [30,100] (corresponding to *r* ∊ [0.0075, 0.025]).

For selective sweeps, we used forward simulations run assuming a diploid population with recombination and mutation parameters as described above. While diploid populations were simulated to incorporate recombination, we used phased haplotypes for our analysis. We assumed a single favored allele under selection coefficient *s* ∊ [0.02, 0.8] and heterozygosity 0.5 (haploid carriers get half the fitness advantage of diploid carriers). When *s* is ‘low’ (0.001 ≤ *s* ≤ 0.01), none of the available tests can detect a selective sweep with reasonable power [21,23]. Selection with *s* ≥ 0.1 is considered ‘high’ (e.g., see [21]). For high values of *s*, the carrier haplotypes are identical or very similar in simulations, making the problem of detecting carriers easy. Therefore, we chose intermediate values (*s* ∊ [0.02, 0.08]) in our simulations.

Soft sweeps can arise either due to standing variation or due to multiple favored alleles. Here, we focus on the former, where the favored allele is present in at least one carrier in the population (*v*_0_ ∊ [1/*N*, 1]), and drifting at the onset of selection. In our simulations, we set *v*_*0*_ at the beginning of the sweep to *v*_*0*_ = 1/*N* for hard sweeps and *v*_0_ = 0.3 for soft sweeps.

### Data Preprocessing

We downloaded pre-phased haplotype data from the HapMap [49] project website. Both HapMap 3 [49] and HapMap 2 [77] project data were used depending on whether the causal allele was sampled or not. For LCT (rs4988235), we used 88 CEU individuals haplotypes from HapMap 3; for PSCA (rs2294008), we used 60 YRI individuals from HapMap 2; for TRPV6, 88 CEU individuals from HapMap 3; for ADH1B (rs1229984), 90 CHB+JPT individuals from HapMap 2; and, for EDAR (rs3827760), we chose 90 CHB+JPT individuals from HapMap 2. The number of phased haplotypes was twice the number of individuals in each case.

We downloaded Chimpanzee genome alignments [50] to identify the ancestral allele. A total of ∼93% of the sites analyzed had were covered by the Chimpanzee data. For these sites, we set the ancestral allele to the Chimpanzee allele, and we discarded sites that were not covered.

## Software

The PreCIOSS software is available from the website http://bix.ucsd.edu/projects/precioss/

## Acknowledgments

This work was supported in part by National Science Foundation grants CCF-1115206 and IIS-1318386.

## Appendix

**Figure S1.**
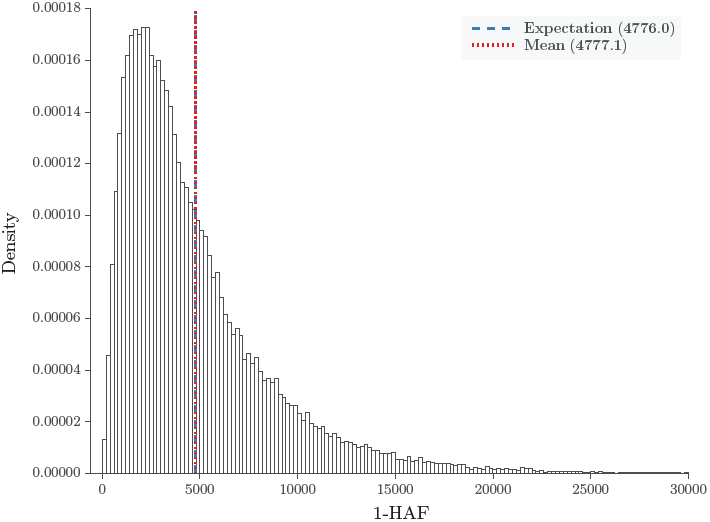
HAF scores in neutrally evolving constant-sized populations. The distribution of 4 × 10^6^ 1-HAF scores aggregated from 20,000 population samples (each of *n* = 200 haplotypes) simulated under a standard coalescent model without recombination. Plugging in the simulation parameters *θ* = 48, *n* = 200, Eqs. (3),(7) evaluate to 4776. The observed mean 1-HAF score is 4777.1.

**Figure S2.**
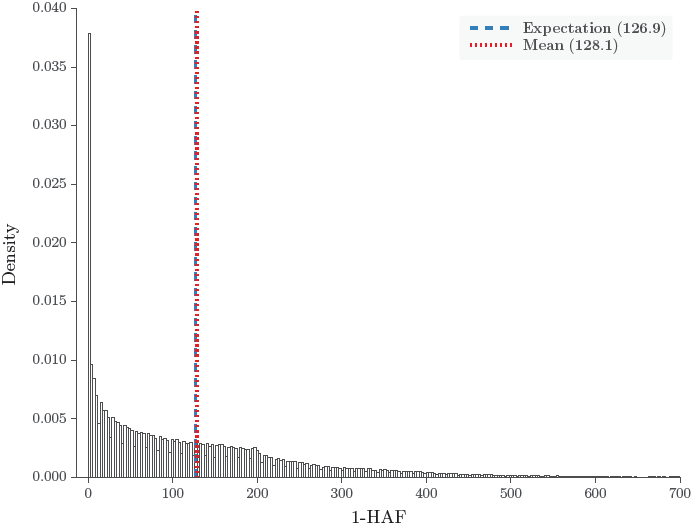
HAF scores in neutrally evolving exponentially growing populations. The distribution of 4 × 10^6^ 1-HAF scores aggregated from 20,000 population samples (each of *n* = 200 haplotypes) simulated under a coalescent model of exponential growth without recombination. Computing the conditional expectation as described in Eq. (12) with the simulation parameters (*θ* = 48, *n* = 200, *α* = 80) gives 126.9. The observed mean 1-HAF score is 128.1.

**Figure S3.**
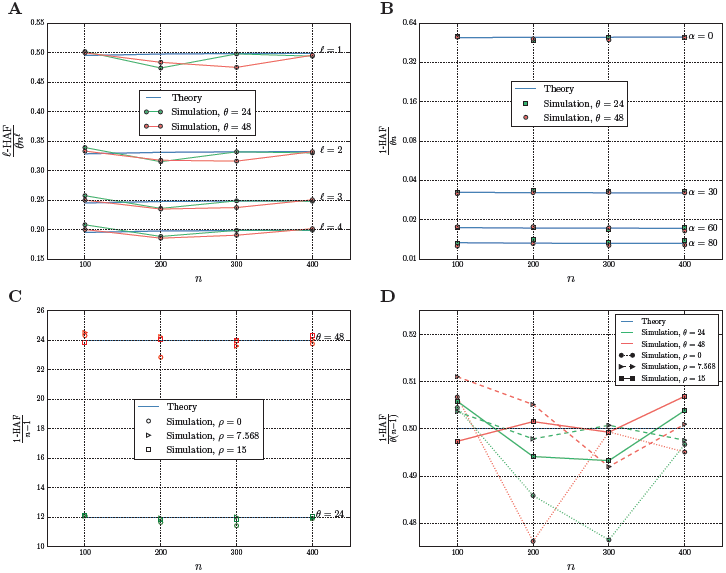
HAF scores for a range of simulation parameters. Each empirical test is the average of 1,000 trials. (A) Empirical mean and theoretical expected 𝓁-HAF scores for a fixed size population (𝓁 ∊ {1, 2, 3, 4}, *θ* ∊ {24,48}, *ρ* = 0). (B) Empirical mean and theoretical expected 1-HAF scores for an exponentially growing population (*α* ∊ {0, 30, 60, 80}, *θ* ∊ {24, 48}, *ρ* = 0). (C) Theoretical expected 1-HAF scores (computed assuming *ρ* = 0) compared against empirical means of 1-HAF scores from samples with different recombination rates (*ρ* ∊ {0,7.568,15}, *θ* ∊ {24,48}). (D) Interestingly, higher recombination rates reduce the variance in 1-HAF estimates. In the three green curves for *θ* = 24 (and in the three red curves for *θ* = 48), the variation from the expected value (blue) decreases as *ρ* increases. Rate *ρ* = 0 (dotted) has the most variation; *ρ* = 7.568 (dashed) has less; and *ρ* = 15 (solid) has the least. The theoretical values are based on (A) Eqs. (3), (S22), (B) Eq. (12), and (C,D) Eq. (4).

**Figure S4.**
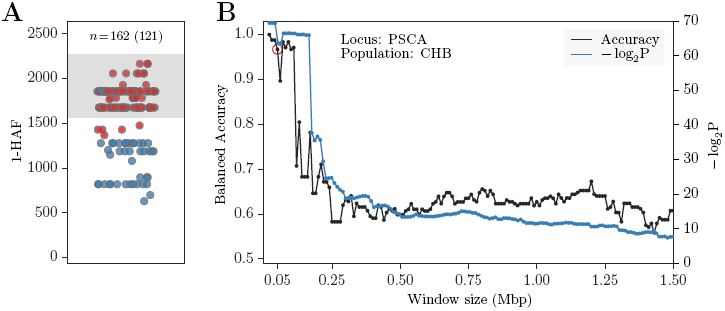
Predicting carriers of the PSCA sweep in CHB. (A) Haplotype 1-HAF scores in a 50 kb window centered at the favored site. (B) Balanced classification accuracy (black) and − log_2_(*P*) values (blue) as function of window size around the favored allele. P-values are for Wilcoxon rank sum tests rejecting the null hypothesis of identically distributed 1-HAF scores among carriers and non-carriers. The red circle indicates the balanced accuracy obtained for the 50 kb window shown on the left. As with the YRI population, we achieve high classification accuracy when considering ∼100 kb window centered at the favored allele. But unlike in YRI, we see a sharp decline in both accuracy and − log_2_(*P*) values beginning at larger distances from the favored allele. See Fig. 6 for further details on the conventions used.

**Figure S5.**
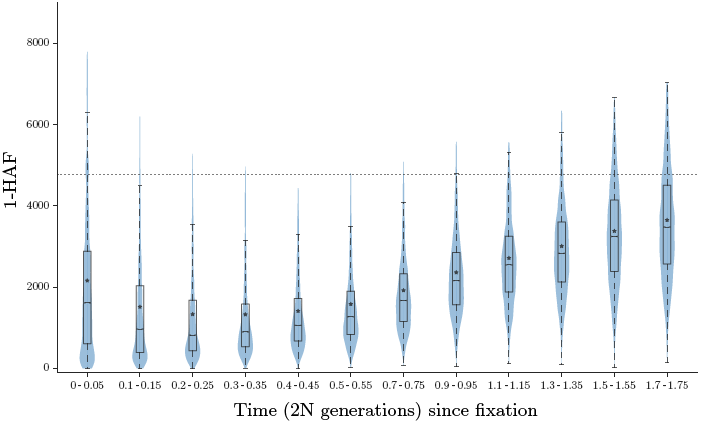
Recovery of HAF scores after a selective sweep. Each violin shows the Gaussian kernel density estimation (KDE) of 1-HAF scores in populations sampled at regular time intervals following the fixation of a selective sweep. All individuals at this stage are carriers of the favored allele. A standard boxplot is overlaid on each violin (outlier points not shown). The horizontal dotted line represents the neutral expected value. HAF scores were computed from 500 simulated populations undergoing a hard sweep, sampled at regular time intervals.

## Empirical validation of PreCIOSS

In Fig. 5A, we showed the balanced accuracy of PreCIOSS for predicting carriers of ongoing hard sweeps. For soft sweeps, the balanced accuracy stays robust but is slightly reduced (Fig. 5B). In Fig. S6 we apply PreCIOSS to data from 500 trials of a simulated population for each sweep type (hard or soft), and for different population genetics parameters provided in Table S1. For soft sweeps, we set *v*_*0*_ = 0.3 (frequency of the favored allele at the onset of selection).

**Table S1.**
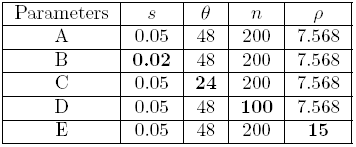
Simulation parameter sets used for generating Fig. S6. In simulations B through E, we changed one parameter (in boldface) at a time vs. simulation A.

| Parameters | $s$ | $\theta$ | $n$ | $\rho$ |
| --- | --- | --- | --- | --- |
| A | 0.05 | 48 | 200 | 7.568 |
| B | **0.02** | 48 | 200 | 7.568 |
| C | 0.05 | **24** | 200 | 7.568 |
| D | 0.05 | 48 | **100** | 7.568 |
| E | 0.05 | 48 | 200 | **15** |

**Figure S6.**
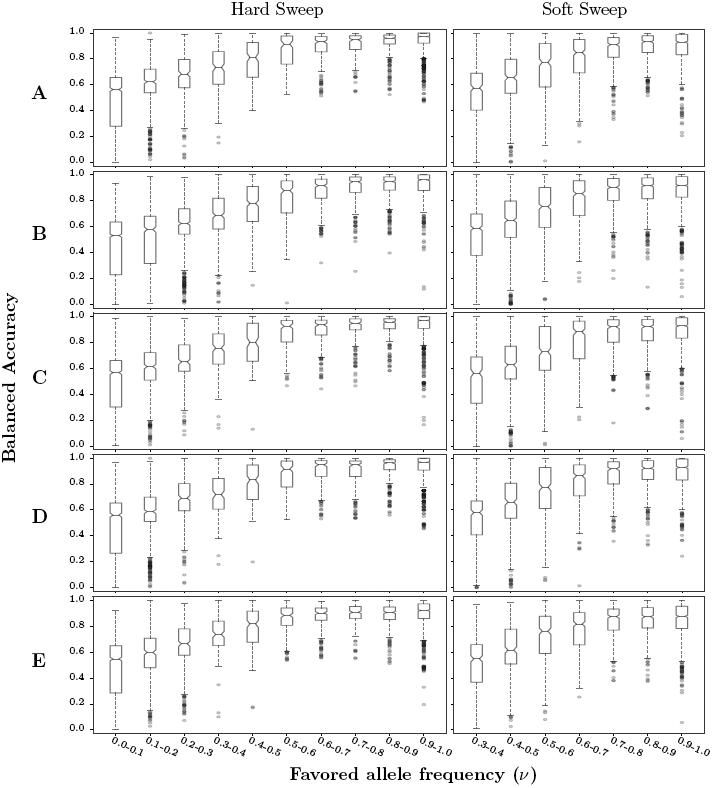
Predicting carriers of hard and soft sweeps. Balanced accuracy of PreCIOSS in populations undergoing hard (left figures) and soft (right figures) sweeps. Balanced accuracy is shown for each allele frequency bin as a standard box plot computed over 500 simulated populations (for each sweep type and each parameter set in Table S1) sampled at regular time intervals during sweep progression.

## Derivation of the expected 𝓁-HAF score

We compute the expected value by averaging over all haplotypes in a genealogy (sum over the haplotypes and divide by *n*) and then averaging over all genealogies. Recall from the main text that *M*_k,i_ and *W*_k,i_ refer to random variables denoting the number of mutations and the frequency, respectively on the *i*^th^ lineage of the *k*^th^ epoch. As the genealogy of a neutrally evolving sample is independent of branch lengths [43], *M*_k,i_ and *W*_k,i_ are independent random variables. Recall Eq. (7):

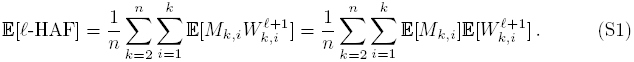

### Expected value of *M*_k,i_, constant population size

Let *T*_*k*_ denote the duration of epoch *k*. For a population of constant size *N*, the duration *T*_*k*_ is exponentially distributed with rate 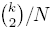 (see [78]):)

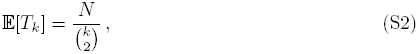

and the number of mutations on a lineage *i* in epoch *k*, denoted *M*_*k,i*_, is Poisson distributed with rate *μT_k_*. For a constant-sized population, this implies

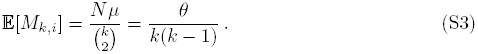

Later, we will explore 𝔼[*T*_*k*_] and 𝔼[*M*_*k,i*_] for exponentially growing populations.

### Rising factorials and moment calculations

The probability distribution of *W*_*k, i*_, or the number of descendants of the *i*^th^ lineage in epoch *k*, is given by [42, Eq. (14)]:

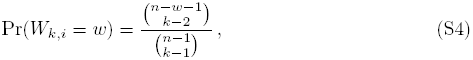

where 1 ≤ *i* ≤ *k*, 2 ≤ *k* ≤ *n*, and 1 ≤ *w* ≤ *n* − *k* + 1. We wish to evaluate Eq. (7) (or (S1)) for 𝓁 = 1, and generally for any 𝓁. This requires 𝔼[*M*_*k,i*_], which we already have for constant populations, and 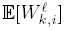, which we derive next. For a positive integer 𝓁, denote the *rising factorial*

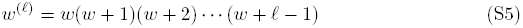

and set *w*^(0)^ = 1. Then 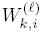 denotes the 𝓁^th^ rising factorial of random variable *W*_*k,i*_, while 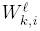 denotes the ordinary 𝓁^th^ power. We will compute 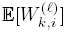, and use it to compute 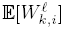.

### Deriving

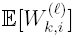

Let 𝓁 be a nonnegative integer. We will prove that

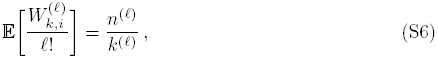

which is equivalent to Eq. (9) in the text. Using Eq. (S4), the expected value is the sum

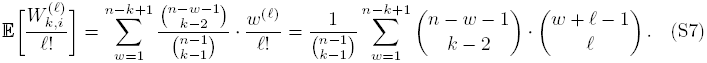

Here we use that rising factorials and binomial coefficients are related via 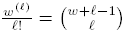

The rightmost sum may be evaluated using a combinatorial method: we will count all (*k* + 𝓁 − 1)-element subsets of {1, 2,…, *n* + 𝓁 − 1} by two methods. First, it is the binomial coefficient 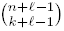. Second, we will show that it also equals the rightmost sum in (S7).

To show this, we may order any such subset and partition it into three parts, (*A, b, C*): *A* is the set of its smallest 𝓁 elements; *b* is the (𝓁 + 1)^th^ smallest element; and *C* is the set of the largest *k* — 2 elements.

Element *b* must be in the range 𝓁 + 1 ≤ *b* ≤ *n* − *k* + 𝓁 + 1 to allow for 𝓁 smaller elements and *k* − 2 larger elements. We rewrite *b* as *b* = 𝓁 + *w*, where *w* is in the range 1 ≤ *w* ≤ *n* − *k* + 1.

Given *b*, we may choose *A* in one of 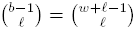 ways and choose *C* in one of 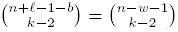 ways. Multiply these counts for a given *b*, and sum over all *b* (by summing over all *w*) to obtain that the total number of (subsets) is the rightmost sum in (S7). But we also showed the number of such subsets is 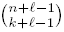. Thus, (S7) evaluates to

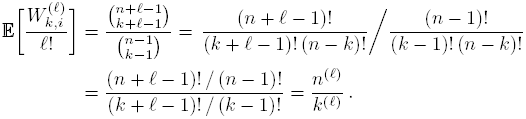

### Deriving 𝔼[𝓁-HAF] in a constant-sized population

Our derivation of 𝔼[𝓁-HAF] consists of three main steps. We start by defining a more general form of the HAF score for an arbitrary function *f* (*w*), which we denote HAF_*f*_. We then set *f*(*w*) to the rising factorial function and derive 𝔼[HAF_*w*^(𝓁)^_]. Finally, we use Stirling Numbers to convert between rising factorials and powers, and obtain 𝔼[𝓁-HAF].

#### Step 1: The generalized form 𝔼[HAF_*f*_]

Consider an arbitrary polynomial *f*(*w*) (or more generally, an arbitrary function *f* : ℤ → ℤ). Define the HAF_*f*_ score of a HAF vector **c** as

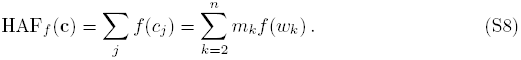

We now generalize the derivation of Eq. (7) to HAF_*f*_. Sum the above equation over all haplotypes in a genealogy:

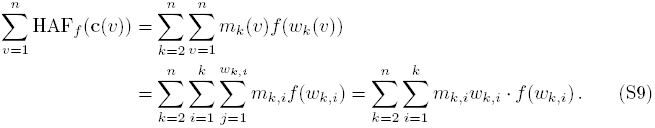

Note that in the sum over *j*, all summands are independent of *j*, so that sum is replaced by multiplying by the number of terms, *w*_*k, i*_. We divide the above sum by *n* and then average over all genealogies to obtain

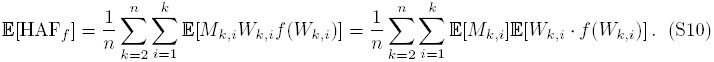

Further, since the random variables *W*_*k,i*_ for *i* = 1,…, *k* are identically distributed, all terms in the inner sum are the same, and may be consolidated into *k* times the value of the term on one branch:

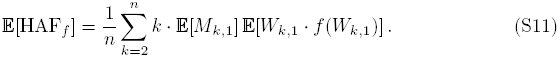

In a constant-sized population, Eq. (S3) states that 𝔼[*M*_*k,i*_] = θ/(*k*(*k* − 1)). Plug this in:

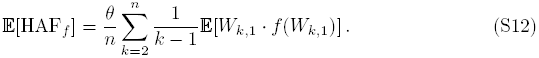

#### Step 2: The rising factorial form 𝔼[HAF_*w*^(𝓁)^_]

We will show that for *f*(*w*) = *w*^(𝓁)^, for any nonnegative integer 𝓁:

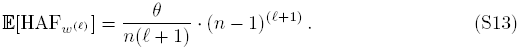

For this proof only, abbreviate *W* = *W*_*k,1*_. Note that

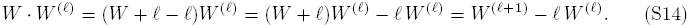

Using this and Eq. (S6) gives

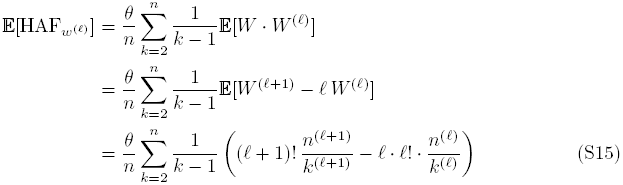

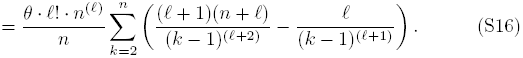

We will split this sum into two telescoping sums. One may show that

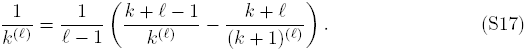

Using this with shifted *k*‘s and 𝓁’s, part of the sum in Eq. (S16) can be written

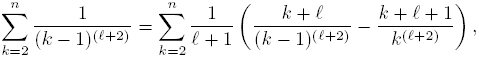

which is a telescoping sum that evaluates to

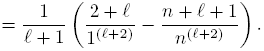

Note that 1^(𝓁+2)^ = (𝓁 + 2)!, simplifying this to

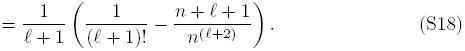

Similarly,

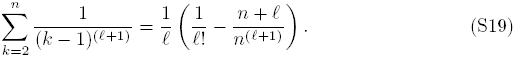

Plugging Eqs. (S18),(S19) into Eq. (S16), we obtain:

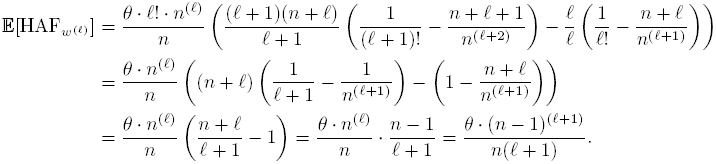

This proves Eq. (S13).

#### Step 3: The final form 𝔼[𝓁-HAF]

Powers *w*^𝓁^ and rising factorials *w*^(𝓁)^ may be expressed as linear combinations of each other via Stirling numbers. Let *c*(*𝓁, q*) denote the Unsigned Stirling Number of the First Kind, and *S*(*𝓁, q*) denote the Stirling Number of the Second Kind. Then [45, p. 264]

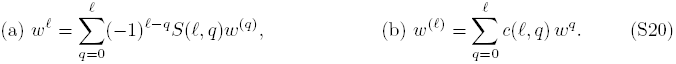

The expected value of (*W*_*k, i*_)^𝓁^ is then:

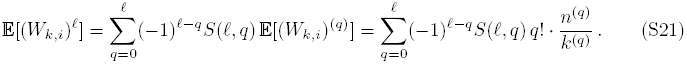

We evaluate 𝔼[𝓁-HAF] = 𝔼[HAF_*w*^(𝓁)^_] as a linear combination of terms 𝔼[HAF_w_(q)], using the same coefficients that express *w*^𝓁^ as a linear combination of terms of *w*^(*q*)^ (Eq. (S20a)):

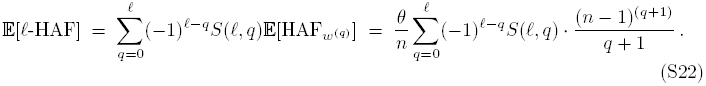

For 𝓁 = 0:

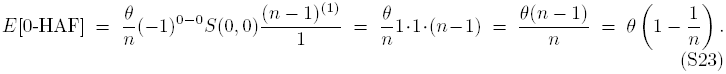

For 𝓁 = 1:

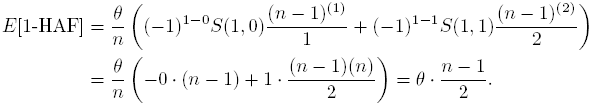

For 𝓁 = 2:

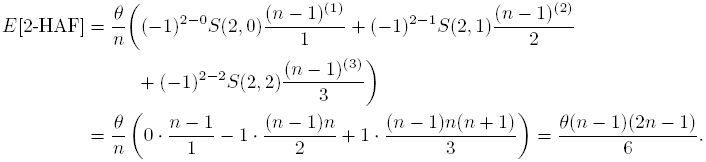

It is straightforward to compute this for any 𝓁 in the same fashion.

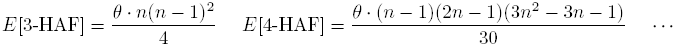

### Equivalence of Eq. (3) and Eq. (7) for 𝔼[𝓁-HAF] in a constant-sized population

Eq. (3) has simple derivation, but a variable number of terms, *n*, that grows as *n* grows. Eq. (7) leads to an evaluation (S22) that is a polynomial in *n* of degree *t*. Both represent 𝔼[t-HAF] in a constant-sized population; we now show that they are equal as a consequence of the following lemma:

#### Lemma 1

*In a constant-sized population, for any polynomial f (w), we have*

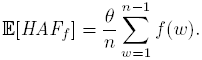

Applying this to *f*(*w*) = *w*^𝓁^ gives equivalence of the two formulas (3) and (S22) for 𝔼[𝓁-HAF].

#### Proof of lemma.

Both sides are linear in *f*, so it suffices to prove it on any basis of polynomials. We use the basis of rising factorials, *f*(*w*) = *w*^(𝓁)^ for 𝓁 ≥ 0. Eq. (S13) evaluates 𝔼[HAF_*f*_] on this basis. We now evaluate 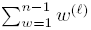. Note that

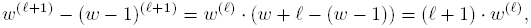

So

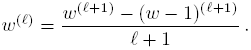

This leads to a telescoping sum:

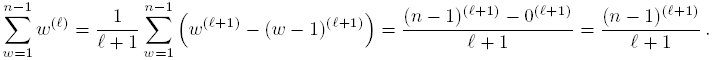

Multiplying this sum by *θ*/n gives the same value as (S13) gives for 𝔼[HAF_*w*_^(𝓁)^]. Thus, the lemma holds on the basis *f*(*w*) = *w*^(𝓁)^ for 𝓁 ≥ 0. By linearity, it holds on all polynomials *f*(*w*).

### Deriving 𝔼[𝓁-HAF] in an exponentially growing population

Let the population size at time *t* in the past be given by *N*(*t*) = *N*_*0*_ *e*^*−rt*^, where *N*_*o*_ is the current population size. *M*_*k,i*_ depends on *T*_*k*_, which in turn depends on the size of the population in epoch *k*. Slatkin and Hudson (1991) [44, p. 559] derived an iterative formula to generate a sequence of random times *T*_*k*_ (*k* = *n*, *n* − 1,…, 2). The actual formula implemented in the simulator *ms* [73] is slightly different; in our notation, it is as follows. Let *α* = 2 *N*_0_ *r* be a scaled growth rate. Generate random values *U*_*n*_,…, *U*_2_ that are uniformly distributed in (0,1), and iteratively compute *T*_*n*_,…, *T*_2_ via

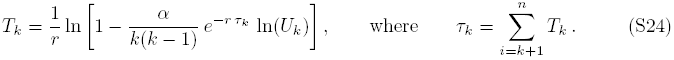

Then generate a random value for each *M*_*k, i*_ using a Poisson distribution with mean *μ*,*T*_*k*_ (*k* = 2,…, *n* and *i* = 1,…, *k*). This does not lead to a closed form expression for so the expected HAF scores, but we may nonetheless use it to estimate the expected HAF scores computationally, as illustrated in Fig. S2.

The first method, *cumulative time,* is to generate random values of *T*_*k*_’s and *M*_*k*_,*i*’s as above, and plug them into (S10), along with exact values of 𝔼[*W*_*k, i*_ · f (*W*_*k, i*_)] computed using (S6) or (S21).

The second method is *conditional expectation.* For a given *k*, we may compute the expected value of (S24) given the condition that *T*_*k*+1_, … , *T*_*n*_ are known. This gives estimates *T*_*k*_ = *t*_*k*_ (computed in order *k* = *n*, *n* − 1,…, 2) as follows:

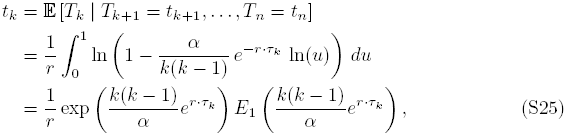

where *τ*_*k*_ = *t*_*k*_+_1_ + · · + *t*_*n*_ (with *τ*_*n*_ = 0) and *E*_*1*_(*x*) is the exponential integral 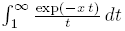.

In practice, Eq. (S25) is used to generate scaled times,

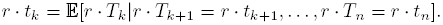

In terms of these times, 𝔼[*M*_*k, i*_] = *μ*𝔼[*T*_*k*_] ≈ *μt_k_* (which is approximate since the formula for *t*_*k*_ uses conditional expectation while this equation does not). We then estimate 𝔼[HAF_*f*_] via (S11):

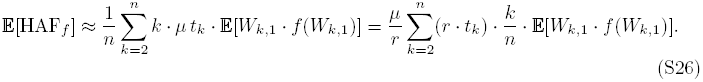

For *f*(*w*) = *w*^(𝓁)^, similarly to (S15), this gives an estimate

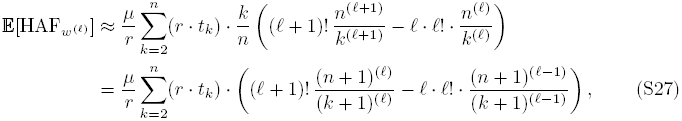

and for *f*(*w*) = *w*^𝓁^, we use (S20) to obtain

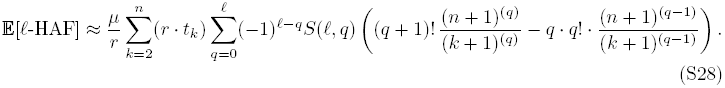

In terms of *α* = 2 *N*_*0*_ *r* and *θ* = 2 *N*_*0*_ *μ*, the coefficient *μ*/*r* may be replaced by *θ*/*α*.

We used Maple (maplesoft.com) to generate the sequence of scaled times *r* · *t*_*k*_ by iterating Eq. (S25). Maple supports arbitrary precision numerical computations, which is needed due to complications with double precision arithmetic. Let 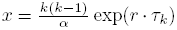. Small rounding errors in x due to limited precision are amplified in e^x^, which may lead to slightly different results when iterating the recursion using double precision arithmetic vs. Maple’s arbitrary precision arithmetic. Additionally, as *x* increases, *e*^*x*^ grows rapidly and leads to an overflow in double precision arithmetic at approximately *x* ≥ 709.7827, while *E*_1_(*x*) decays rapidly and leads to an underflow at approximately *x* ≥ 701.8334, even though *e*^*x*^*E*_1_(*x*) is usually representable in double precision. For example, *E*_1_(702) ≈ 1.9 · 10^−308^ underflows in double precision arithmetic, but *e*^702^*E*_1_(702) ≈ 0.0014 is representable. The largest value of *x* used in Eq. (S25) is *x* = *n*(*n* − 1)/*α* in iteration *k* = *n*, leading to an underflow in double precision if *n*(*n* − 1)/*α* > 701.8.

### Number of mutations in a genealogy

Let *f*(*w*) = 1/*w* for *w* > 0 and *f*(0) = 0. Note that for an individual HAF vector **c**, some entries may be 0, but in a genealogy, all *w*_*k, i*_ ≠ 0 since every branch of the observed tree has at least one haplotype as its descendant. The sum (S9) of HAF_*f*_ (**c**) over all haplotypes in a genealogy becomes

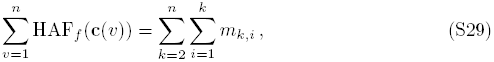

and thus (S11) becomes

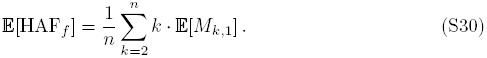

In a constant-sized population, Eqs. (S3) and (S29) give that mutations in a genealogy is

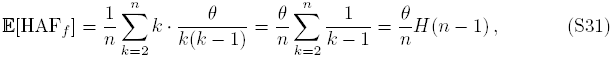

where 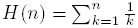 is the Harmonic number.

Under exponential growth, we instead estimate times *t*_*k*_ or scaled times *r* · *t*_*k*_ via (11). Then we estimate the expected number of mutations by

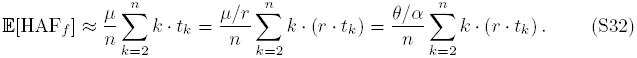

### Computing the mean HAF_*f*_ score from a SNP matrix

For a sample of *n* individuals whose genealogy has *q* segregating sites, the SNP matrix *A* consists of n rows (each row representing one individual) and *q* columns, with entry 0 denoting ancestral alleles and 1 denoting derived alleles.

The *v*^th^ row of A is a haplotype vector as depicted as **h** in Fig. 1.

Let **w** ^a11^ denote the sum of the rows of *A*. The frequency of the *j*^th^ allele is 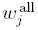, the sum of all entries in the *j*^th^ column of *A*.

In 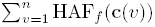, the *j*^th^ mutation contributes *f*(*w*_*j*_) for each of the *w*_*j*_ haplotypes that have that mutation (*A*_*vj*_ = 1) and contributes 0 on each of the *n* − *w*_*j*_ others. Thus, the average of HAF_*f*_ over the genealogy is

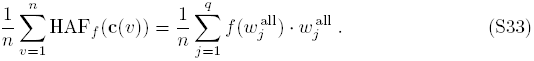

We use this to compute empirical averages of HAF_*f*_ in simulations (*ms* [73] outputs a SNP matrix rather than a tree), as well as in further theoretical developments in this Appendix. In particular, given a haplotype vector **h** and the corresponding HAF vector **c**, we have

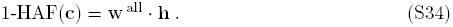

### HAF score dynamics and peak values

Consider *n* haplotypes randomly sampled from a Wright-Fischer (WF) fixed-size population of *N* haploid individuals under a hard sweep with selection coefficient s. Let v denote the fraction of individuals that carry the favored allele. We assume the sample has at least one carrier and at least one non-carrier, so that 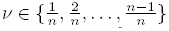 at fixation (*v* = 1), there are no non-carriers remaining. We assume strong selection (*N*_*s*_ ≫ 1) and no recombination in the region being sampled. We measure time in generations going backwards. See Fig. S7. The current time is time 0; let *T*^car^ denote the time when all sampled carriers coalesce to their most recent common ancestor (denoted by MRCA^car^). Let *T*^a11^ denote the time when all sampled individuals coalesce to a common ancestor (denoted by MRCA^a11^). Let *T*(*k*) denote time to MRCA of *k* randomly chosen haplotypes in a population of size *N*. From Nordborg [47], we have

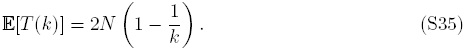

At time *T*^car^, we have exactly one ancestor of the favored allele carrier, and assume we have *m* ancestors of non-carriers. In a hard-sweep scenario, the favored mutation arises at the same time as the onset of selection; therefore, the time between *T*^car^ and *T*^a11^ is governed by the neutral WF model with population *N*, and is well approximated by neutral coalescent theory. Applying Eq. (S35) to the remaining sample of *m* + 1 individuals at time *T*^car^ gives the expected time to coalesce:

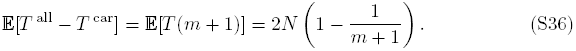

Let *A* denote the SNP matrix of a sample of *n* individuals, with *vn* carriers of the favored allele (Fig. S8). We order the columns so that all mutations are ordered chronologically from left to right. Similarly, the rows are ordered so that the first *vn* rows correspond to carrier haplotypes.

The *i*^th^ haplotype is represented by the *i*^th^ row of *A* (denoted by row vector **h_i_**) and has 1-HAF score denoted by 1-HAF_i_. From Eq. (S34), we have

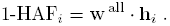

We partition the matrix into 3 submatrices: *A*_1_ consists of the rows corresponding to carriers, and columns corresponding to mutations occurring prior to the onset of selection (times between *T*^car^ and *T* ^a11^). *A*_2_ is the submatrix from all non-carrier rows, and mutations prior to onset of selection. Finally, *A*_3_ is the submatrix of all columns after the onset of selection (times between 0 and *T*^car^). Define *w*^car^ as the sum of all carrier haplotypes; similarly, let *w*^non^ be the sum of all non-carrier haplotypes. Thus,

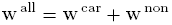

Also, we can partition the equation for computing 1-HAF_i_ by rows as

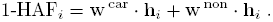

We partition a haplotype vector **h** in the matrix by separating SNPs that occurred before (subvector **h**^b^) and after (subvector **h**^a^) the onset of mutation. A similar partitioning works for **w**^a11^, **w**^car^, and **w**^non^. Thus, for a random haplotype **h**,

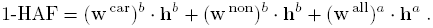

The expected 1-HAF score of a carrier (or non-carrier) haplotype h can similarly be decomposed as

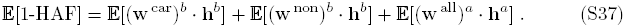

To compute the expected 1-HAF score, we bound each of these constituent terms.

**Figure S7.**
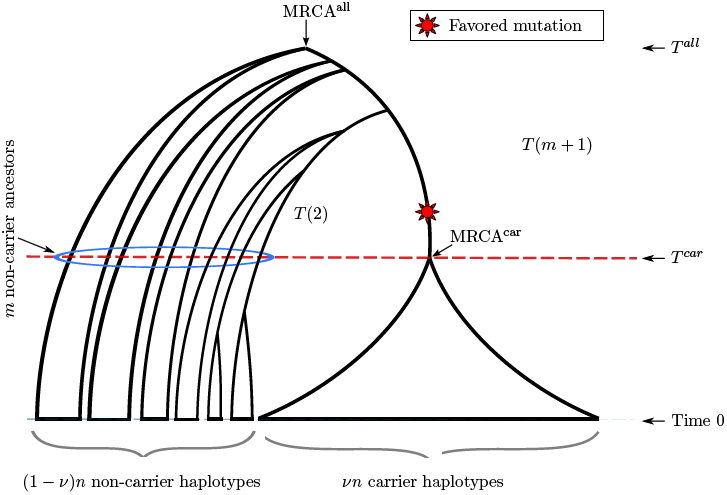
The coalescence of a sample of *n* individuals to their most recent common ancestor MRCA^a11^, during a hard sweep. We assume that the current time has *vn* carriers of the favored allele. These coalesce to MRCA^car^ in *T*^car^ generations. From that point, the coalescence to MRCA^a11^ is governed by neutral coalescent theory. T(k) is time to MRCA of *k* randomly chosen haplotypes in a neutrally evolving population.

**Figure S8.**
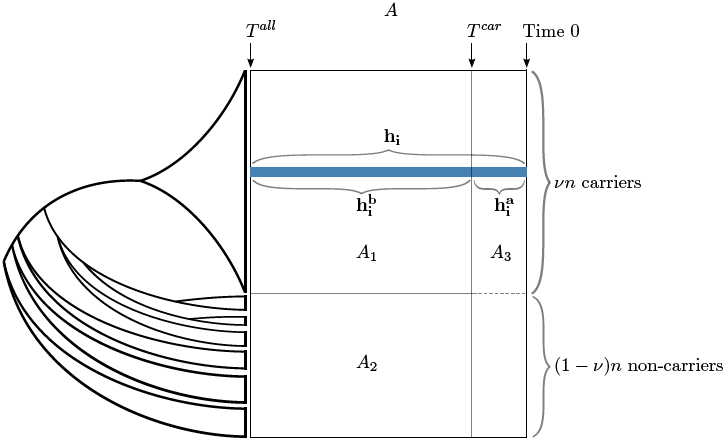
Partitioning the SNP matrix A of a sample of *n* individuals.

#### Lemma 2

Consider a sample of n individuals under a hard sweep with vn carriers 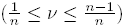. Let *m* be the number of non-carrier haplotypes remaining at time *T*^car^. Then for a random carrier haplotype **h**, we have

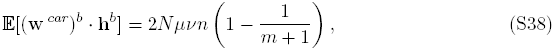

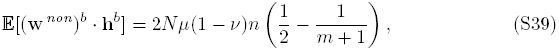

while for a random non-carrier haplotype **h**, these become

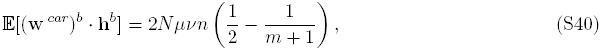

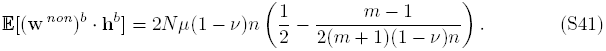

#### Proof of Eq. (S38).

Mutations that happen on the lineage from MRCA^a11^ to MRCA^car^ are shared by all carriers, so all rows of matrix *A*_1_ are identical. Thus, for all carrier haplotypes **h**_*i*_, **h**_*j*_, we have 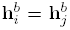. On restricting 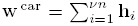 to the mutations from before the onset of selection, we obtain

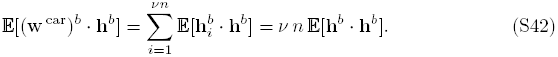

The term 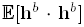 is the expected number of ones in **h**^*b*^, which is the expected number of mutations on the lineage from MRCA^a11^ to MRCA^car^. Therefore, 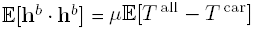. By Eq. (S36),

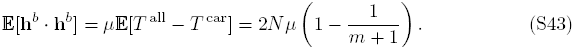

Plugging this into Eq. (S42) gives Eq. (S38). ▀

#### Proof of Eq. (S39).

On restricting 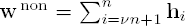 to the mutations from before the onset of selection, we obtain

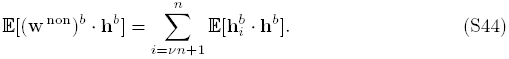

Going back from time *T*^car^ to *T*^a11^, we observe the coalescence of the ancestors of carrier haplotype **h** and non-carrier haplotype **h**_*i*_ at a time *T*′, where *T*^car^ ≤ *T*′ ≤ *T*^a11^. The term 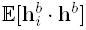 is the expected number of ones in common between these two haplotypes among the SNPs occurring before the onset of selection. This equals the expected number of mutations in the lineage from *T*′ to *T*^a11^:

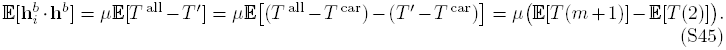

Note that *T*′ — *T*^car^ is the time for a sample of 2 individuals (ancestors of **h** and **h**_*i*_ at time *T*^car^) to coalesce. We evaluate Eq. (S45) using Eq. (S35):

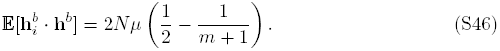

Plug this into Eq. (S44). All *n* − *vn* = (1 − *v*)*n* terms are the same, so it simplifies to Eq. (S39).▀

#### Proof of Eq. (S40).

We have

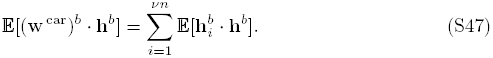

Here, **h** is a non-carrier while **h**_*i*_ is a carrier, so they are distinct and they coalesce prior to the onset of selection. Thus, Eq. (S46) applies here. All *vn* terms of the sum in Eq. (S47) evaluate to Eq. (S46), so it simplifies to Eq. (S40).▀

#### Proof of Eq. (S41).

We have

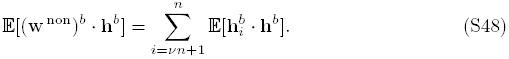

The term 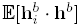 is the expected number of ones in common in haplotypes **h** and **h**_*i*_ among the SNPs occurring before the onset of selection.

We apply Lemma 3 (upcoming) to show that two random non-carrier haplotypes **h** and **h**_i_ have the same ancestor at time *T*^car^ with probability

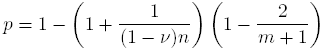

or different ancestors with probability 1 − *p*. Thus, with probability *p*, we have 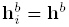, and with probability 1 − *p*, we have 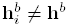. Therefore,

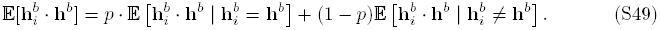

Term 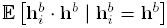 is the expected number of ones in **h**^*b*^, which is 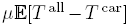. Term 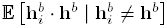 is the expected number of ones in common between 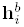 and **h**^*b*^, which is 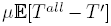. These expected times were evaluated in Eqs. (S43) and (S45). Then

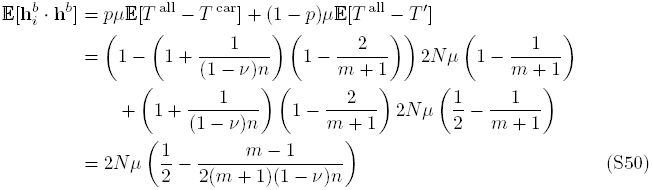

Eq. (S48) is a sum of (1 − *v*)*n* terms, each of form (S50), so Eq. (S48) evaluates to Eq. (S41).▀

In the preceding proof, the probability *p* that two non-carriers share the same ancestor was computed using the following lemma, applied to *n*_0_ = (1 − *v*)*n* non-carriers in the sample and *m* ancestors of non-carriers at time *T*^car^.

#### Lemma 3

Consider a subsample of n_0_ individuals, which coalesce to m ancestors at some time T, while the rest of the sample coalesces to one or more ancestors distinct from these m. Draw two individuals (with replacement) from the subsample. The probability that they have the same ancestor at time T is

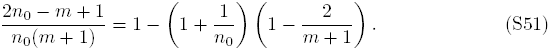

#### Proof.

Throughout this proof, pairs of individuals (*x*, *y*) are drawn from the subsample of size *n*_0_ *with replacement.* There are 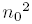 such pairs.

Consider the m lineages at time *T*. The number of pairs of individuals deriving from the *i*^th^ lineage is (*w*_*m,i*_)^2^. Sum over all lineages to obtain the total number of pairs of individuals with the same time *T* ancestor: 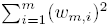. The expected number of pairs with the same ancestor is

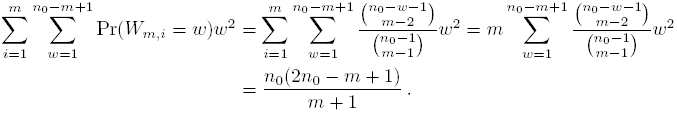

Note that Pr(*W*_*m,i*_ = *w*) is given by Eq. (S4), and holds upon restricting the full genealogy to the subsample. Divide this expected value by 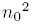 pairs and simplify to obtain Eq. (S51).▀

#### Lemma 4

Consider a sample of *n* individuals under a hard sweep. For a random haplotype **h** (carrier or non-carrier), we have

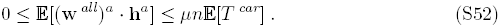

#### Proof.

We have

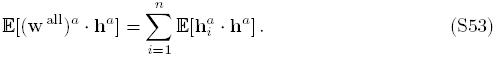

The dimension of vectors 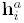 and **h** is the number of mutations in [0, *T*^car^]. The expected value of this dimension is *μ***E**[*T*^car^]. The dot product of two binary vectors is at least 0, and is bounded above by their dimension, so

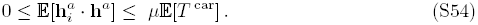

Plugging Eq. (S54) into Eq. (S53) gives Eq. (S52). ▀

#### Lemma 5

Consider a population of size *N* with selection coefficient s, under a hard sweep with strong selection (*N*_*s*_ ≫ 1). Then

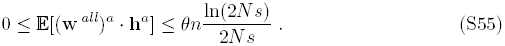

#### Proof.

Campbell [79] shows that under strong selection, the fixation time is approximately 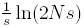. As *T*^car^ is the time for the favored allele to reach frequency *vn*, it is dominated by the fixation time. Therefore,

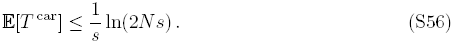

Plug Eq. (S56) and the scaled mutation rate *θ* = 2*Nμ* into Lemma 4 to obtain Eq. (S55). ▀

#### Lemma 6

Consider a sample of n individuals under a hard sweep with vn carriers 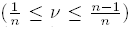. In the time [0, *T*^car^], let the number of coalescent events among the (1 − *v*)*n* non-carrier haplotypes be *ε*_*v*_(1 − *v*)*n*. Then

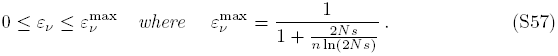

#### Proof.

Since we have (1 − *v*)*n* non-carrier haplotypes in our sample, the expected number of non-carriers in the whole population is (1 − *v*)*N*. Let 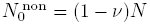, and let 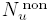 be the expected number of non-carriers in the whole population when the *u*^th^ coalescent event occurs among the non-carrier samples. As we go backwards in time (increasing *u*), the expected number of non-carriers increases:

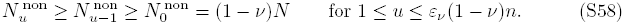

Additionally, the chance of coalescing decreases and the expected coalescent time increases. For a fixed size population of size *N*, the expected time for one coalescent event in a sample of size *n* is 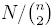 (see [47]). Hence, the expected time for the *u*^th^ coalescent event is at least

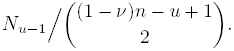

On the other hand, the sum of all coalescent times during the selective sweep is at most *T*^car^. Therefore,

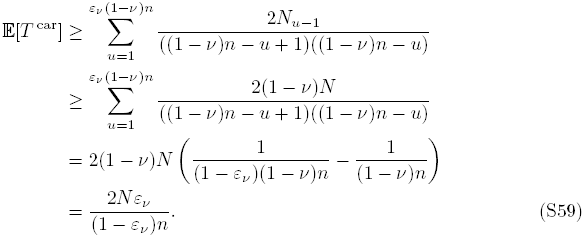

Comparing this lower bound on 𝔼[*T*^car^] with the upper bound in Eq. (S56) gives

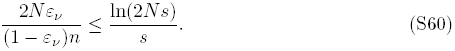

This gives the upper bound on *ε*_*v*_ in Eq. (S57). ▀

#### Theorem 7

Consider a sample of *n* individuals under a hard sweep with *vn* carriers 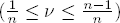. Assume strong selection (*N*_*s*_ ≫ 1). In the time [0, *T*^car^], let the number of coalescent events among the (1 − *v*)*n* non-carrier haplotypes be *ε*_v_(1 − *v*)*n*. The expected 1-HAF score of a random carrier haplotype is

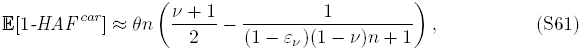

while for a random non-carrier haplotype,

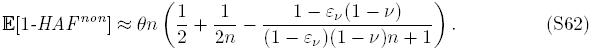

#### Proof.

Recall the decomposition of 𝔼[1-HAF] given in Eq. (S37):

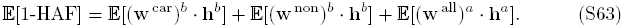

The number of ancestors of non-carriers at time *T*^car^ is *m* = (1 − *ε*_*v*_)(1 − *v*)*n*.

For a random carrier haplotype **h**, we evaluate the first two terms in Eq. (S63) by using Eqs. (S38) and (S39), and bound the third term by using Lemma 5, to obtain:

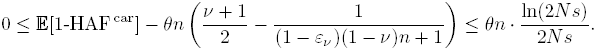

For a random non-carrier haplotype h, we use Eqs. (S40) and (S41) and Lemma 5:

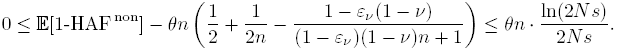

Ignoring the small term 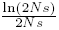, we obtain the desired expressions (S61) and (S62).▀

We find the maximum of Eq. (S61) over 0 ≤ *v* ≤ 1 using differentiation; the result is

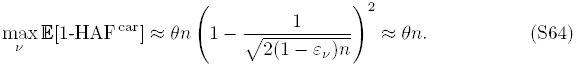

We simulated selective sweeps for a variety of parameters and compared their trajectories against these results. In Fig. S9, we compared the trajectories of both carriers and non-carriers in 500 selective sweeps for each pair (*θ*, *n*) with *θ* ∊ {24, 48}, *n* ∊ {100, 200, 300, 400}, *s* = 0.08, and *n* = 2000. Over the course of a trajectory, the frequency *v* of the favored allele varies from *v* = 1/*n* (1 carrier) to *v* = 1 (fixation), but different trajectories will go through a different sequence of values of *v*. We aligned the trajectories by the values of *v*. For each *v*, we separated carriers and non-carriers and averaged together (1-HAF)/(*nθ*) over the generation (if any) with allele frequency *v* in each sweep. We plotted these averaged trajectories in green (*θ* = 24) and pink (*θ* = 48). We compared these against the expected value (Theorem 7). The expected value is bounded above in blue (*ε*_*v*_ = 0) and below in red 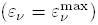; see Lemma 6. These are bounds on the expected value, not on simulation means. Simulation means may range over the whole distribution, but tend to vary around the expected value.

In Fig. S10, for each (*n*, *θ*), we compute average simulated values of (1-HAF)/*θ* over 500 trajectories, stratified by *v* by the same method. The maximum value of this average over all *v* is determined and plotted, and compared against the expected value (Eq. (S64)).

In this paper, due to the run time for forward simulations, we used a relatively small value for population size (*N* = 2000); in reality, *N* is a lot larger (10,000), so that *ε*_*v*_ becomes smaller (see Lemma 6). Therefore, the upper and lower bounds in the proof of Theorem 7 move closer together, and we approximate *ε*_*v*_ = 0. To simplify the presentation in the main text, we set *ε*_*v*_ = 0, giving:

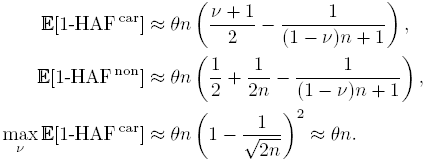

**Figure S9.**
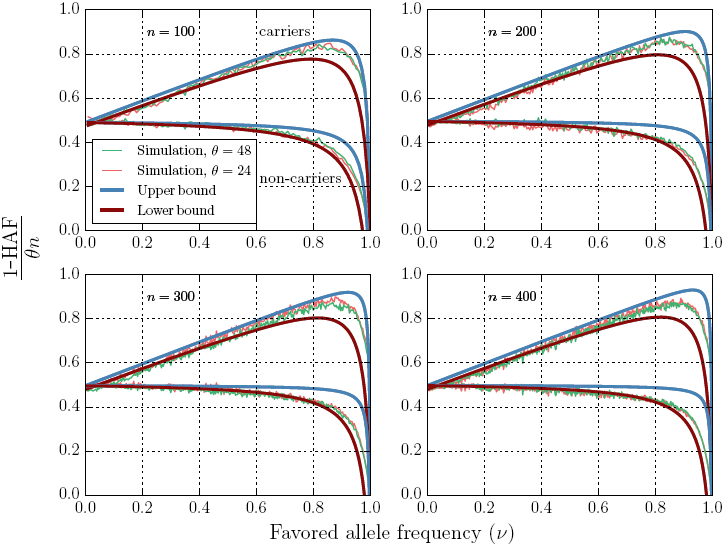
Dynamics of expected 1-HAF score during a selective sweep. For each (*θ*,*n*,*v*) with *θ* ∊ {24,48}, *n* ∊ {100, 200, 300, 400}, 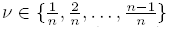, _s_ = 0.08, and *N* = 2000, we did 1500 trials. We plotted the mean value of (1-HAF)/(*θ*n) as a function of v, for both carriers and non-carriers, and compared against the theoretical expected value. The expected value of (1-HAF)/(*nθ*) lies somewhere between the blue and red curves. The mean values may range over the whole distribution (and are not constrained by the blue and red curves) but tend to vary around the expected value.

**Figure S10.**
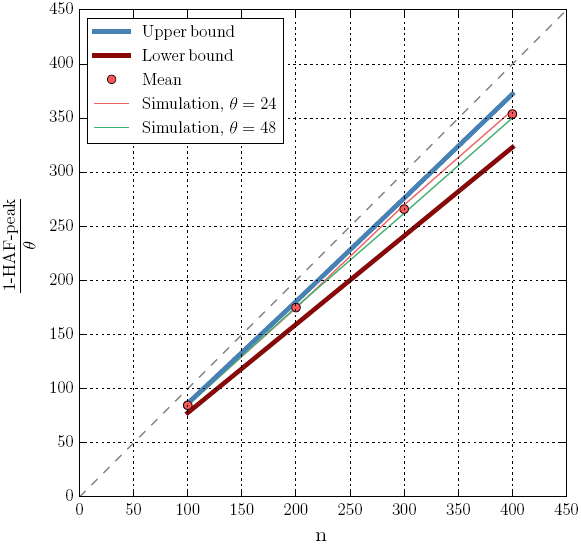
HAF-peak as a function of (*θ, n*. For each (*θ*, *n*, *v*) with *θ* ∊ {24, 48}, *n* ∊ {100, 200, 300, 400}, and 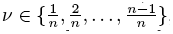, we did 1500 trials. We averaged the values of (1-HAF)/*θ* for carriers at each *v* across the sweeps and determined the maximum over all *v*. Each thin curve shows the results for a fixed *θ* as *n* varies, while the thick curves in red and blue give the theoretical lower and upper bounds on 𝔼[1-HAF ^car^]/*θ*. The solid red spots are the average over all trials over all *θ*, for each *n*.

## References

1. Fu W, Akey JM. Selection and adaptation in the human genome. Annu Rev Genomics Hum Genet. 2013;14:467–489.

2. Lachance J, Tishkoff SA. Population Genomics of Human Adaptation. Annu Rev Ecol Evol Syst. 2013 Nov;44:123–143.

3. Vitti JJ, Grossman SR, Sabeti PC. Detecting natural selection in genomic data. Annu Rev Genet. 2013;47:97–120.

4. Nielsen R, Williamson S, Kim Y, Hubisz MJ, Clark AG, Bustamante C. Genomic scans for selective sweeps using SNP data. Genome Res. 2005 Nov;15(11):1566–1575.

5. Pickrell JK, Coop G, Novembre J, Kudaravalli S, Li JZ, Absher D, et al. Signals of recent positive selection in a worldwide sample of human populations. Genome Res. 2009 May;19(5):826–837.

6. Chen H, Patterson N, Reich D. Population differentiation as a test for selective sweeps. Genome Res. 2010 Mar;20(3):393–402.

7. Berg JJ, Coop G. A population genetic signal of polygenic adaptation. PLoS Genet. 2014 Aug;10(8):e1004412.

8. Jeong C, Di Rienzo A. Adaptations to local environments in modern human populations. Curr Opin Genet Dev. 2014 Dec;29C:1–8.

9. Tekola-Ayele F, Adeyemo A, Chen G, Hailu E, Aseffa A, Davey G, et al. Novel genomic signals of recent selection in an Ethiopian population. Eur J Hum Genet. 2014 Nov; advance online publication.

10. Yi X, Liang Y, Huerta-Sanchez E, Jin X, Cuo ZXP, Pool JE, et al. Sequencing of 50 Human Exomes Reveals Adaptation to High Altitude. Science. 2010;329(5987):75–78. Available from: http://www.sciencemag.org/content/329/5987/75.abstract.

11. Simonson TS, Yang Y, Huff CD, Yun H, Qin G, Witherspoon DJ, et al. Genetic evidence for high-altitude adaptation in Tibet. Science. 2010 Jul;329(5987):72–75.

12. Scheinfeldt LB, Soi S, Thompson S, Ranciaro A, Woldemeskel D, Beggs W, et al. Genetic adaptation to high altitude in the Ethiopian highlands. Genome Biol. 2012;13(1):R1.

13. Alkorta-Aranburu G, Beall CM, Witonsky DB, Gebremedhin A, Pritchard JK, Di Rienzo A. The genetic architecture of adaptations to high altitude in Ethiopia. PLoS Genet. 2012;8(12):e1003110.

14. Huerta-Sanchez E, Degiorgio M, Pagani L, Tarekegn A, Ekong R, Antao T, et al. Genetic signatures reveal high-altitude adaptation in a set of ethiopian populations. Mol Biol Evol. 2013 Aug;30(8):1877–1888.

15. Udpa N, Ronen R, Zhou D, Liang J, Stobdan T, Appenzeller O, et al. Whole genome sequencing of Ethiopian highlanders reveals conserved hypoxia tolerance genes. Genome Biol. 2014 Feb;15(2):R36.

16. Zhou D, Udpa N, Ronen R, Stobdan T, Liang J, Appenzeller O, et al. Whole-genome sequencing uncovers the genetic basis of chronic mountain sickness in Andean highlanders. Am J Hum Genet. 2013 Sep;93(3):452–462.

17. Kaplan NL, Hudson RR, Langley CH. The “hitchhiking effect” revisited. Genetics. 1989 Dec;123(4):887–899.

18. Smith JM, Haigh J. The hitch-hiking effect of a favourable gene. Genet Res. 1974 Feb;23(1):23–35.

19. Tajima F. Statistical method for testing the neutral mutation hypothesis by DNA polymorphism. Genetics. 1989 Nov;123(3):585–595.

20. Fay JC, Wu CI. Hitchhiking under positive Darwinian selection. Genetics. 2000 Jul;155:1405–1413.

21. Pavlidis P, Jensen JD, Stephan W. Searching for footprints of positive selection in whole-genome SNP data from nonequilibrium populations. Genetics. 2010 Jul;185(3):907–922.

22. Lin K, Li H, Schlotterer C, Futschik A. Distinguishing positive selection from neutral evolution: boosting the performance of summary statistics. Genetics. 2011 Jan;187(1):229–244.

23. Ronen R, Udpa N, Halperin E, Bafna V. Learning natural selection from the site frequency spectrum. Genetics. 2013 Sep;195(1):181–193.

24. Simonsen KL, Churchill GA, Aquadro CF. Properties of statistical tests of neutrality for DNA polymorphism data. Genetics. 1995 Sep;141(1):413–429.

25. Braverman JM, Hudson RR, Kaplan NL, Langley CH, Stephan W. The hitchhiking effect on the site frequency spectrum of DNA polymorphisms. Genetics. 1995 Jun;140(2):783–796.

26. Hudson RR, Bailey K, Skarecky D, Kwiatowski J, Ayala FJ. Evidence for positive selection in the superoxide dismutase (Sod) region of Drosophila melanogaster. Genetics. 1994 Apr;136(4):1329–1340.

27. Depaulis F, Mousset S, Veuille M. Haplotype tests using coalescent simulations conditional on the number of segregating sites. Mol Biol Evol. 2001 Jun;18(6):1136–1138.

28. Innan H, Zhang K, Marjoram P, Tavare S, Rosenberg NA. Statistical tests of the coalescent model based on the haplotype frequency distribution and the number of segregating sites. Genetics. 2005 Mar;169(3):1763–1777.

29. Sabeti PC, Reich DE, Higgins JM, Levine HZ, Richter DJ, Schaffner SF, et al. Detecting recent positive selection in the human genome from haplotype structure. Nature. 2002 Oct;419(6909):832–837.

30. Voight BF, Kudaravalli S, Wen X, Pritchard JK. A map of recent positive selection in the human genome. PLoS Biol. 2006 Mar;4(3):e72.

31. Toomajian C, Hu TT, Aranzana MJ, Lister C, Tang C, Zheng H, et al. A nonparametric test reveals selection for rapid flowering in the Arabidopsis genome. PLoS Biol. 2006 May;4(5):e137.

32. Sabeti PC, Varilly P, Fry B, Lohmueller J, Hostetter E, Cotsapas C, et al. Genome-wide detection and characterization of positive selection in human populations. Nature. 2007 Oct;449(7164):913–918.

33. Kim Y, Stephan W. Selective sweeps in the presence of interference among partially linked loci. Genetics. 2003 May;164(1):389–398.

34. Messer PW, Petrov DA. Population genomics of rapid adaptation by soft selective sweeps. Trends Ecol Evol (Amst). 2013 Nov;28(11):659–669.

35. Hermisson J, Pennings PS. Soft sweeps: molecular population genetics of adaptation from standing genetic variation. Genetics. 2005 Apr;169(4):2335–2352.

36. Pennings PS, Hermisson J. Soft sweeps II–molecular population genetics of adaptation from recurrent mutation or migration. Mol Biol Evol. 2006 May;23(5):1076– 1084.

37. Ferrer-Admetlla A, Liang M, Korneliussen T, Nielsen R. On detecting incomplete soft or hard selective sweeps using haplotype structure. Mol Biol Evol. 2014 May;31(5):1275–1291.

38. Garud NR, Messer PW, Buzbas EO, Petrov DA. Recent selective sweeps in North American Drosophila melanogaster show signatures of soft sweeps. PLoS Genet. 2015 Feb;11(2):e1005004.

39. Peter BM, Huerta-Sanchez E, Nielsen R. Distinguishing between selective sweeps from standing variation and from a de novo mutation. PLoS Genet. 2012;8(10):e1003011.

40. Schrider DR, Mendes FK, Hahn MW, Kern AD. Soft Shoulders Ahead: Spurious Signatures of Soft and Partial Selective Sweeps Result from Linked Hard Sweeps. Genetics. 2015 Feb; advance online publication.

41. Wilson BA, Petrov DA, Messer PW. Soft selective sweeps in complex demographic scenarios. Genetics. 2014 Oct;198(2):669–684.

42. Fu YX. Statistical properties of segregating sites. Theor Popul Biol. 1995 Oct;48(2):172–197.

43. Hudson RR. Gene genealogies and the coalescent process. In: Futuyma D, Antonovics J, editors. Oxford Surveys in Evolutionary Biology. Oxford: Oxford University Press; 1990. p. 1–44.

44. Slatkin M, Hudson RR. Pairwise comparisons of mitochondrial DNA sequences in stable and exponentially growing populations. Genetics. 1991 Oct;129(2):555–562.

45. Graham R, Knuth DE, Patashnik O. Concrete Mathematics: A Foundation for Computer Science. 2nd ed. Reading, Mass: Addison-Wesley; 1994.

46. Durrett R. Probability Models for DNA Sequence Evolution. 2nd ed. Probability and Its Applications. Springer; 2008.

47. Nordborg M. Coalescent Theory. In: Balding DJ, Bishop M, Cannings C, editors. Handbook of statistical genetics. 3rd ed. John Wiley & Sons, Ltd; 2008. p. 843–877.

48. Brodersen KH, Ong CS, Stephan KE, Buhmann JM. The Balanced Accuracy and Its Posterior Distribution. In: Pattern Recognition (ICPR), 2010 20th International Conference on; 2010. p. 3121–3124.

49. Altshuler DM, et al. Integrating common and rare genetic variation in diverse human populations. Nature. 2010 Sep;467(7311):52–58.

50. Sequencing TC, Consortium A. Initial sequence of the chimpanzee genome and comparison with the human genome. Nature. 2005 Sep;437(7055):69–87.

51. Kuokkanen M, Enattah NS, Oksanen A, Savilahti E, Orpana A, Jarvela I. Transcriptional regulation of the lactase-phlorizin hydrolase gene by polymorphisms associated with adult-type hypolactasia. Gut. 2003 May;52(5):647–652.

52. Olds LC, Sibley E. Lactase persistence DNA variant enhances lactase promoter activity in vitro: functional role as a cis regulatory element. Hum Mol Genet. 2003 Sep;12(18):2333–2340.

53. Troelsen JT, Olsen J, Møller J, Sjöström H. An upstream polymorphism associated with lactase persistence has increased enhancer activity. Gastroenterology. 2003 Dec;125(6):1686–1694.

54. Akey JM, Eberle MA, Rieder MJ, Carlson CS, Shriver MD, Nickerson DA, et al. Population history and natural selection shape patterns of genetic variation in 132 genes. PLoS Biol. 2004 Oct;2(10):e286.

55. Stajich JE, Hahn MW. Disentangling the effects of demography and selection in human history. Mol Biol Evol. 2005 Jan;22(1):63–73.

56. Akey JM, Swanson WJ, Madeoy J, Eberle M, Shriver MD. TRPV6 exhibits unusual patterns of polymorphism and divergence in worldwide populations. Hum Mol Genet. 2006 Jul;15(13):2106–2113.

57. Bhatia G, Patterson N, Pasaniuc B, Zaitlen N, Genovese G, Pollack S, et al. Genome-wide comparison of African-ancestry populations from CARe and other cohorts reveals signals of natural selection. Am J Hum Genet. 2011 Sep;89(3):368–381.

58. Sakamoto H, Yoshimura K, Saeki N, Katai H, Shimoda T, Matsuno Y, et al. Genetic variation in PSCA is associated with susceptibility to diffuse-type gastric cancer. Nat Genet. 2008 Jun;40(6):730–740.

59. Wu X, Ye Y, Kiemeney LA, Sulem P, Rafnar T, Matullo G, et al. Genetic variation in the prostate stem cell antigen gene PSCA confers susceptibility to urinary bladder cancer. Nat Genet. 2009 Sep;41(9):991–995.

60. Whitfield JB. Alcohol dehydrogenase and alcohol dependence: variation in genotype-associated risk between populations. Am J Hum Genet. 2002 Nov;71(5):1247–1250.

61. Peng Y, Shi H, Qi XB, Xiao CJ, Zhong H, Ma RL, et al. The ADH1B Arg47His polymorphism in east Asian populations and expansion of rice domestication in history. BMC Evol Biol. 2010;10:15.

62. Osier MV, Pakstis AJ, Soodyall H, Comas D, Goldman D, Odunsi A, et al. A global perspective on genetic variation at the ADH genes reveals unusual patterns of linkage disequilibrium and diversity. Am J Hum Genet. 2002 Jul;71(1):84–99.

63. Eng MY, Luczak SE, Wall TL. ALDH2, ADH1B, and ADH1C genotypes in Asians: a literature review. Alcohol Res Health. 2007;30(1):22–27.

64. Li H, Mukherjee N, Soundararajan U, Tarnok Z, Barta C, Khaliq S, et al. Geographically separate increases in the frequency of the derived ADH1B*47His allele in eastern and western Asia. Am J Hum Genet. 2007 Oct;81(4):842–846.

65. McGovern PE, Zhang J, Tang J, Zhang Z, Hall GR, Moreau RA, et al. Fermented beverages of pre- and proto-historic China. Proc Natl Acad Sci USA. 2004 Dec;101(51):17593–17598.

66. Fujimoto A, Ohashi J, Nishida N, Miyagawa T, Morishita Y, Tsunoda T, et al. A replication study confirmed the EDAR gene to be a major contributor to population differentiation regarding head hair thickness in Asia. Hum Genet. 2008 Sep;124(2):179–185.

67. Kimura R, Yamaguchi T, Takeda M, Kondo O, Toma T, Haneji K, et al. A common variation in EDAR is a genetic determinant of shovel-shaped incisors. Am J Hum Genet. 2009 Oct;85(4):528–535.

68. Bryk J, Hardouin E, Pugach I, Hughes D, Strotmann R, Stoneking M, et al. Positive selection in East Asians for an EDAR allele that enhances NF-kappaB activation. PLoS ONE. 2008;3(5):e2209.

69. Sabeti PC, Varilly P, Fry B, Lohmueller J, Hostetter E, Cotsapas C, et al. Genome-wide detection and characterization of positive selection in human populations. Nature. 2007 Oct;449(7164):913–918.

70. Williamson SH, Hernandez R, Fledel-Alon A, Zhu L, Nielsen R, Bustamante CD. Simultaneous inference of selection and population growth from patterns of variation in the human genome. Proc Natl Acad Sci USA. 2005 May;102(22):7882–7887.

71. Luksza M, Lassig M. A predictive fitness model for influenza. Nature. 2014 Mar;507(7490):57–61.

72. Lee MC, Lopez-Diaz FJ, Khan SY, Tariq MA, Dayn Y, Vaske CJ, et al. Single-cell analyses of transcriptional heterogeneity during drug tolerance transition in cancer cells by RNA sequencing. Proc Natl Acad Sci USA. 2014 Nov;111(44):E4726–4735.

73. Hudson RR. Generating samples under a Wright-Fisher neutral model of genetic variation. Bioinformatics. 2002 Feb;18(2):337–338.

74. Nachman MW, Crowell SL. Estimate of the mutation rate per nucleotide in humans. Genetics. 2000 Sep;156(1):297–304.

75. Campbell CD, Chong JX, Malig M, Ko A, Dumont BL, Han L, et al. Estimating the human mutation rate using autozygosity in a founder population. Nat Genet. 2012 Nov;44(11):1277–1281.

76. Fledel-Alon A, Leffler EM, Guan Y, Stephens M, Coop G, Przeworski M. Variation in human recombination rates and its genetic determinants. PLoS ONE. 2011;6(6):e20321.

77. Frazer KA, et al. A second generation human haplotype map of over 3.1 million SNPs. Nature. 2007 Oct;449(7164):851–861.

78. Kingman JFC. On the genealogy of large populations. Journal of Applied Probability. 1982;19:27–43.

79. Campbell RB. Coalescent size versus coalescent time with strong selection. Bull Math Biol. 2007 Oct;69(7):2249–2259.

